# An Omics approach on *Marchantia polymorpha* single FERONIA and MARIS homologs confirms links between cell wall integrity and abscisic acid

**DOI:** 10.1101/2024.11.26.625412

**Authors:** Timothy Owen Jobe, Celso Gaspar Litholdo, Sara Christina Stolze, Lisa Stephan, Jens Westermann, Anne Harzen, Martin Hülskamp, Hirofumi Nakagami, Aurélien Boisson-Dernier

**Affiliations:** Institute for Plant Sciences, University of Cologne, 50674 Cologne, Germany; U.S. Department of Agriculture, Agriculture Research Service, Plant Stress and Water Conservation Laboratory, 3810 4th Street, Lubbock, TX 79415, USA.; Université Côte d’Azur, INRAE, CNRS, Institut Sophia Agrobiotech, 06903 Sophia Antipolis Cedex, France; Max Planck Institute for Plant Breeding Research, Cologne, Germany

**Keywords:** *Marchantia polymorpha*, Malectin-like Receptor Kinases, FERONIA, MARIS, Cell Wall Integrity, RNA-Seq, LC-MS/MS, Abscisic Acid, GSE282327, PXD058089

## Abstract

Plant cells are surrounded by an extracellular cell wall that shields them from their abiotic and biotic environment. To coordinate their growth with their cell wall status, plant cells have developed cell wall integrity (CWI) mechanisms, at the center of which lies the transmembrane Malectin-like receptor kinase FERONIA (FER). FER controls a myriad of plant developmental processes including sexual reproduction, cell growth and morphogenesis, often intersecting with phytohormones-dependent pathways such as abscisic acid (ABA) signaling or plant immunity. Interestingly, FER together with its downstream receptor-like cytoplasmic kinase MARIS (MRI) was shown to similarly control root hair and rhizoid integrity in the vascular angiosperm Arabidopsis and the early diverging bryophyte *Marchantia polymorpha*, respectively.

Here, we performed comparative transcriptomics and proteomics on the *M. polymorpha* mutant plants, Mp*fer-1* and Mp*mri-1,* and their corresponding wild-type accessions Tak-1 and Tak-2. Large and significant overlaps were observed between differentially expressed genes and differentially abundant proteins in both mutants. Our multi-omics approach revealed that MpFER and MpMRI largely cooperate to negatively regulate transcriptional and translational networks, particularly those related to plant defense and ABA responses. Moreover, our phenotypic analyses showed that Mp*fer-1* plants are hypersensitive to ABA-dependent growth inhibition, indicating that FER’s function of negatively regulating ABA-related growth responses is conserved between bryophytes and vascular plants.

## INTRODUCTION

In the last 30 years, studies on the plant cell wall (CW), the extracellular matrix surrounding plant cells and preventing them from bursting, have revealed that this extracellular compartment is much more than a rigid frame. Rather, the CW is an extremely dynamic and complex biochemical barrier that adapts and communicates actively with the cell protoplast it shelters (Cosgrove, 2024; Delmer *et al*., 2024). Part of this communication is arbitrated by plasma membrane localized receptor-like proteins and receptor-like kinases whose extracellular domains perceive and bind signals within the CW matrix to regulate intracellular activities (Wolf, 2022). One of these receptor classes is the Malectin-like receptor-kinase (MLR) family also known as the CrRLK1L family, with 17 members in the flowering plant model Arabidopsis (Franck *et al*., 2018; Zhu *et al*., 2021). MLRs contain an intracellular Ser/Thr kinase, a transmembrane domain and an extracellular-domain with 2 malectin-like domains that can bind pectins of the CW and small secreted peptides called RALFs (Rapid Alkalinization Factors). FERONIA (FER) is by far the most studied of the MLRs: it is expressed widely throughout the plant body -except in pollen where its pollen-expressed sister genes ANXUR1 and 2, and BUDDHA’S PAPER SEALl and 2 are taking over (Miyazaki *et al*., 2009; Boisson-Dernier *et al*., 2009; Ge *et al*., 2017)- and is involved in a myriad of biological processes and signaling pathways (Malivert & Hamant, 2023; Cheung, 2024). FER was originally discovered to function in the female gametophyte as a key component for the dialogue between male and female gametophytes that results in sperm release in the synergids and hence was named after the Etruscan goddess of fertility (Rotman *et al*., 2003; Huck *et al*., 2003). Since then, FER, together with its co-receptor LLG1 (LORELEI-LIKE glycosylphosphatidylinositol-anchored protein) and their peptide ligand RALFs, has been shown to orchestrate many developmental processes including root (Haruta *et al*., 2014) and root hair growth (Duan *et al*., 2010), as well as pavement cell morphogenesis (Li *et al*., 2015; Lin *et al*., 2022). The involvement of FER in such processes operates at the crossroad with many phytohormone signaling pathways such as Brassinosteroids (Guo *et al*., 2009), Ethylene (Deslauriers & Larsen, 2010), and Abscisic Acid (ABA; Yu *et al*., 2012; Chen *et al*., 2016) and in conjunction with external stress perception such as high salinity (Feng *et al*., 2018), and pathogen invasions (Kessler *et al*., 2010; Masachis *et al*., 2016; Stegmann *et al*., 2017). Roots of mutant plants for FER and LLG1 carry root hairs that burst prematurely during growth (Duan *et al*., 2010; Li *et al*., 2015). Similarly, pollen tubes of double mutants for ANX1/ANX2 and their co-receptor LLG2/LLG3 burst *in vitro* and *in vivo* (Miyazaki *et al*., 2009; Boisson-Dernier *et al*., 2009; Ge *et al*., 2017; Feng *et al*., 2019). These studies demonstrate that MLRs and their co-receptors are required to maintain cell wall integrity (CWI) during tip-growth. Interestingly, a receptor-like cytoplasmic kinase from the RLCK-VIII subfamily named MARIS (MRI), after the Etruscan god of fertility and agriculture, was identified as a common downstream regulator of these signaling receptor complexes (Boisson-Dernier *et al*., 2015). *MRI* is preferentially expressed in both pollen tubes and root hairs of Arabidopsis and *mri* mutants also display pollen and root hair bursting (Boisson-Dernier *et al*., 2015; Liao *et al*., 2016). Moreover, expression of an overactive variant of *MRI*, *MRI^R240C^*, is sufficient to rescue the bursting phenotype of *fer* root hairs and *anx1 anx2* pollen tubes, positioning MRI downstream of the MLRs (Boisson-Dernier *et al*., 2015). The R240C mutation in the kinase activation loop of MRI is likely protecting MRI from the regulatory inhibition of a yet unknown protein phosphatase 2C as it was reported for the *Nicotiana benthamiana* MRI homolog Pti1b (Pto interacting 1b; (Giska & Martin, 2019)). While FER and MRI have not yet been reported to interact or phosphorylate each other, MRI was recently shown to be phosphorylated *in vitro* by BUPS1 while ANX1 failed to do so (Gao *et al*., 2023b). In addition, MRI and MRI^R240C^ were found to interact with pollen-expressed MILDEW LOCUS O1 (MLO1), and overactive MRI^R240C^, but not MRI, could activate the calcium channel activity of AtMLO1, 5, 9 and 15 (Gao *et al*., 2023b). Altogether, these studies indicate that members of the RALF/LLG/MLR/MRI/MLO families constitute a signaling module that regulates cell wall integrity during tip growth. Excitingly, this signaling module active in pollen tubes and root hairs of the flowering plant Arabidopsis was also found to be functional in the rhizoids of the bryophyte *Marchantia polymorpha* (hereafter referred to as *Marchantia*; Westermann *et al*., 2019). Rhizoids are tip-growing cells that participate in anchoring the plant body to the substrate and absorbing water and nutrients, and as such, are considered analogous structures of root hairs (Menand *et al*., 2007). T-DNA insertional knock-down mutants for Mp*FER* and Mp*MRI*, the unique homologues of the MLR and RLCK-VIII families in *Marchantia* respectively, displayed bursting rhizoids while expression of Mp*MRI^R240C^* was sufficient to partially rescue Mp*fer-1* bursting rhizoids (Westermann *et al*., 2019). Thus, the MLR/MRI signaling module that controls CWI in tip-growing cells has been conserved throughout land plant evolution from the rhizoids of bryophytes to the root hairs and pollen tubes of seed plants. In addition, transspecies complementation assays showed that MpMRI and AtMRI (64% identity at the amino acid level) can functionally replace each other in the control of tip growth (Westermann *et al*., 2019), while this was not true for MpFER and AtFER (43.3% identity at the amino acid level; Mecchia *et al*., 2022). Thus, it remains uncertain which of the known MLR and MRI functions in Arabidopsis originate from mechanisms established early during land plant evolution and which ones have arisen in the last 450 million years ago.

Here, we exploit the knock-down mutants of the CWI module in the liverwort *Marchantia* by carrying out comparative transcriptomic (RNA-seq) and proteomic (LC-MS/MS) analyses of Mp*fer-1* and Mp*mri-1* compared to the wild-type accessions Tak-1 and Tak-2. Our results show that MpFER and MpMRI largely cooperate to negatively regulate transcriptional and translational networks, in particular those related to plant defense and abscisic acid (ABA) responses. Moreover, Mp*fer-1* displayed hypersensitivity to ABA-induced thalli growth inhibition, indicating that the negative regulation of ABA signaling by MpFER and AtFER is conserved and active in the common ancestor of land plants.

## MATERIALS AND METHODS

### Marchantia material and growth conditions

The *Marchantia polymorpha* Tak-1 and Tak-2 ecotypes as well as the Mp*fer-1* and Mp*mri-1* T-DNA insertional mutant lines (Honkanen *et al*., 2016; Westermann *et al*., 2019) were cultivated via propagation of vegetative propagules (gemmae) on solid Johnson’s medium (Ishizaki *et al*., 2008) supplemented with 0.8 % micro agar under axenic conditions. Cultivation petri dishes were sealed using Micropore tape to ensure gas exchange while preventing microbe contamination. Gemmae were grown under long day conditions (16 h light/8 h darkness cycle) and white light irradiation (60 µmol m-1 s-1) at 21°C and 60 % humidity.

For ABA-growth inhibition assays, fresh gemmae from cups were placed on square Petri dishes containing agar solidified ½ strength Gamborg’s B5 medium, pH 5.7, complemented or not with 1 or 10 μM ABA (A1049; Sigma-Aldrich) for 14 days.

### Transcriptomic Analyses

Total RNA for RNA-seq analysis was extracted from four biological replicates of 3-week-old Tak-1, Mp*fer-1*, Tak-2 and Mp*mri-1* thalli grown on cellophane membranes to preserve the rhizoids using TRI reagent (Ambion Life Technologies). Two μg of DNaseI-treated, RNA (RIN > 7, OD260/280 = 1.8–2.1, OD260/230 > 1.5) for three of the four biological replicates were sent to the Cologne Center for Genomics (CCG) for paired-end 100 bp short-read sequencing. Raw sequencing data was imported into the Galaxy web server and the read quality was assessed using the *FastQC* package. Adapter sequences were trimmed, and the reads were quality-filtered using the *fastp* package with default settings and enabling overrepresentation analysis. Transcript count was then determined from the resulting preprocessed FASTQ files using the *KallistoQuant* package with default settings and sequence-based bias correction (42 bootstraps). Finally, differentially expressed genes (DEGs) were determined using the *DESeq2* package with default settings. An adjusted *p*-value ≤ 0.05 and log_2_(fold change) ≥1.5 or ≤-1.5 were set as the cutoff values to identify differentially expressed transcripts. All transcripts were mapped to the *Marchantia polymorpha* genome (MpTakv6.1) obtained from the MarpolBase website (https://marchantia.info). The RNA-seq data from this study are available at the NCBI Gene Expression Omnibus (GEO) repository under accession number GSE282327.

To visualize the robustness of our experimental design and assess the quality of the biological replicates used in our RNA-seq experiment, we performed several analyses. First, PCA plots were generated from the transcript counts of each individual replicate using the *plotPCA* package in Galaxy (Figure 1B). Next, we extracted a list of DEGs and the corresponding transcript abundances from each replicate. To visualize these data, heatmaps were generated using R Studio with the ggplot2 and pheatmap packages.

**Figure 1:**
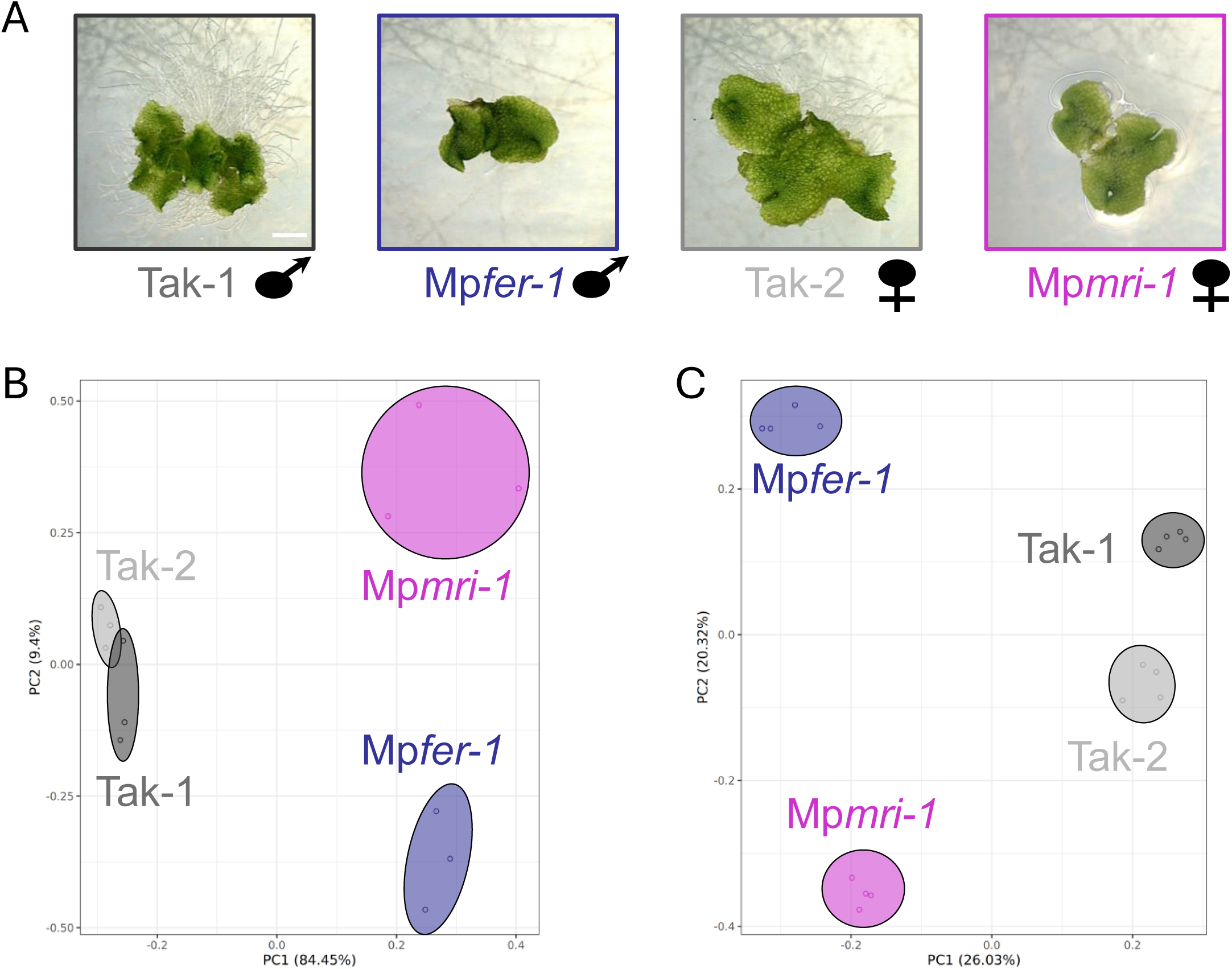
Comparative omics studies of the *Marchantia* cell wall integrity (CWI) mutants. A) Examples of thalli and rhizoid growth on cellophane membranes of the wild-types (WT) Tak-1 (male, dark grey) and Tak-2 (female, light grey), and the CWI mutants Mp*fer-1* (male, dark purple) and Mp*mri-1* (female, light purple). Thalli and their rhizoids were used for transcriptomic and proteomic analyses. B) Principal component analysis of RNA-seq data obtained from three biological replicates of the Tak-1, Tak-2, Mp*fer-1*, and Mp*mri-1* genotypes. C) Principal component analysis of proteomic data obtained from four biological replicates of the Tak-1, Tak-2, Mp*fer-1*, and Mp*mri-1* genotypes.

Heatmaps were generated for each wild-type/mutant pair (Mp*fer-1* vs Tak-1, and Mp*mri-1* vs Tak-2). Using the same list of differentially expressed transcripts, we then plotted the log_2_(fold change) vs the -log_10_(p-value) with R studio and the ggplot2 package to generate volcano plots showing the magnitude and significance of differentially expressed transcripts for Tak-2 vs Tak-1 (Figure S1A) and each wild-type/mutant pair (Figure 2A, and 3A).

**Figure 2:**
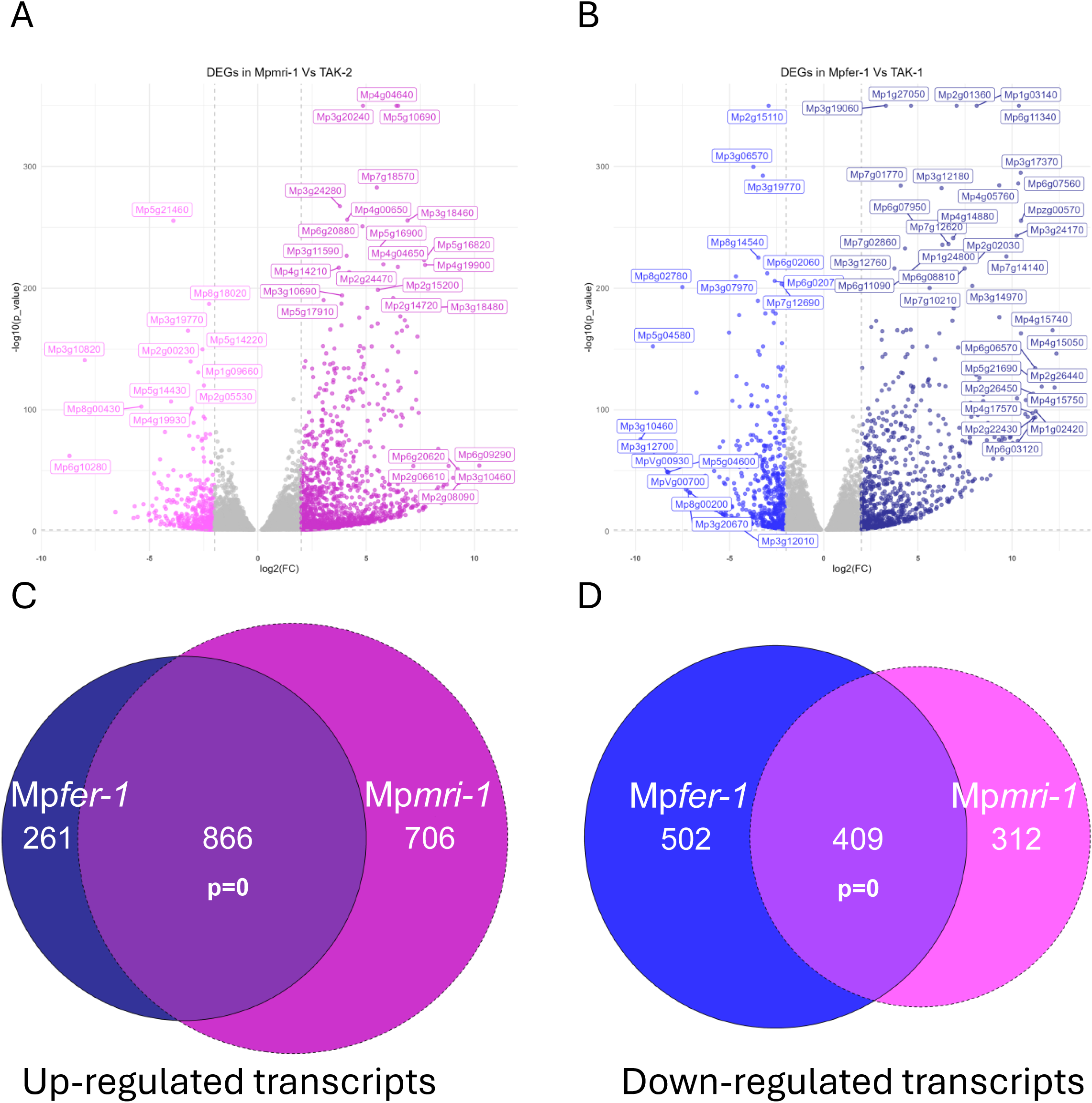
Volcano plots and Venn diagrams showing differential abundance of transcripts in the *Marchantia* CWI mutants and their overlaps. A) Transcriptional analysis showing DEGs that are up (dark pink) or down (light pink) regulated in Mp*mri-1* compared to Tak-2. B) Transcriptional analysis showing DEGs that are up (dark blue) or down (light blue) regulated in Mp*fer-1* compared to Tak-1. Differential expression was determined based on log_2_ (FC) ≥ 1.5 and p-value ≤ 0.05. C) Venn diagram showing the total number of up-regulated DEGs in Mp*fer-1* and Mp*mri-1* and the number of DEGs common to both analyses. D) Venn diagram showing the total number of down-regulated DEGs in Mp*fer-1* and Mp*mri-1* and the number of DEGs common to both analyses. The statistical significance of overlapping datasets in C) and D) was determined using a hypergeometric test.

Next, we determined the number of common and unique transcripts by comparing the list of differentially expressed transcripts in Mp*fer-1* vs Tak-1 to those in Mp*mri-1* vs Tak-2. To determine the significance of overlapping transcriptomic datasets we used the hypergeometric test from the R *phyper* function and visualized the overlap using area-proportional Euler diagrams with EulerR (Figure 2C, D, Figure 3C, D, Figure S2, S8).

**Figure 3:**
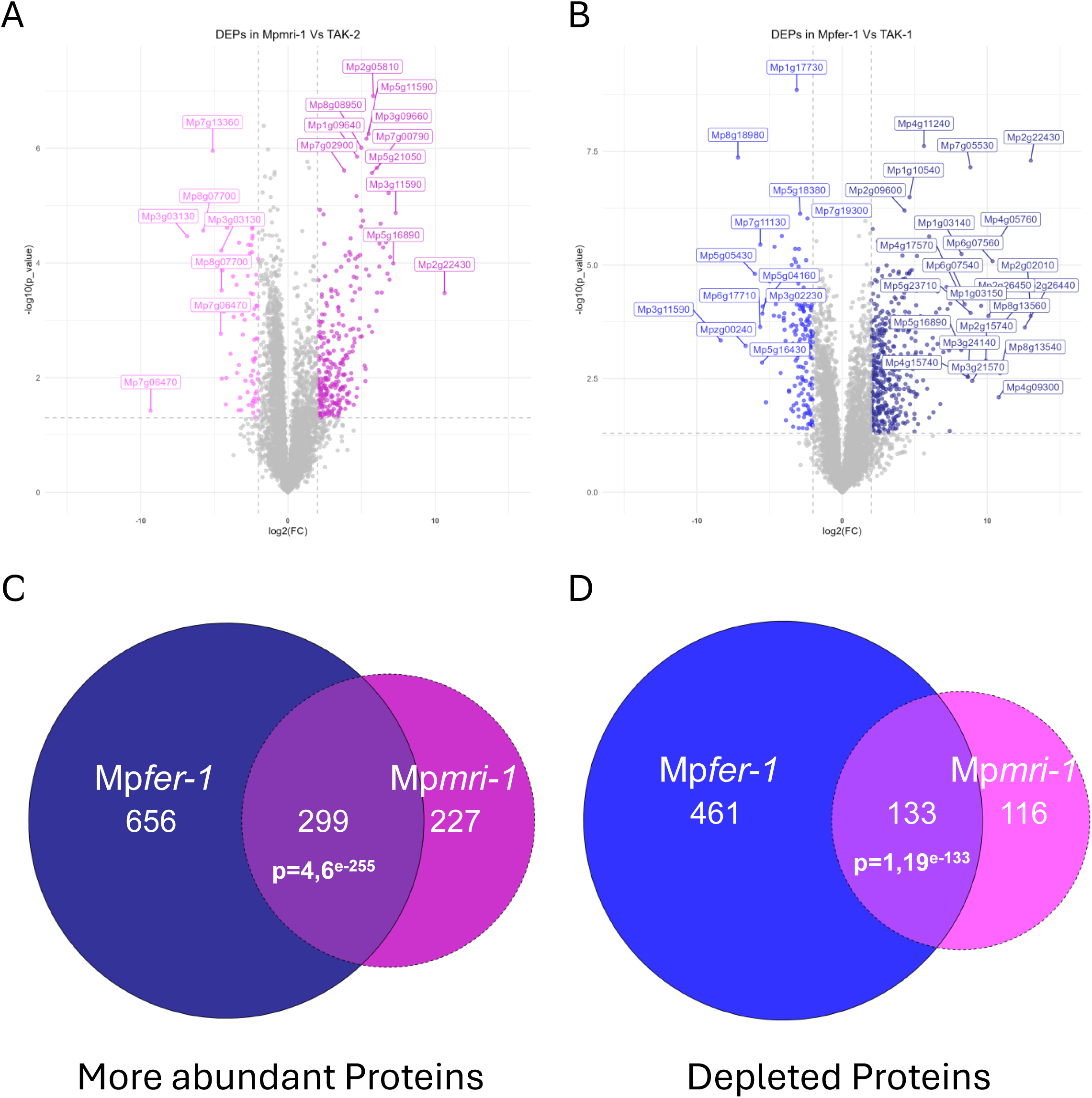
Volcano plots and Venn diagrams showing differential abundance of proteins in the *Marchantia* CWI mutants and their overlaps. A) Proteomic analysis showing DEPs that are up (dark pink) or down (light pink) regulated in Mp*mri-1* compared to Tak-2. B) Transcriptional analysis showing DEPs that are up (dark blue) or down (light blue) regulated in Mp*fer-1* compared to Tak-1. C) Venn diagram showing the total number of up-regulated DEPs in Mp*fer-1* and Mp*mri-1* and the number of DEPs common to both analyses. Differential abundance was determined based on log_2_ (FC) ≥ 2 and p-value ≤ 0.05. D) Venn diagram showing the total number of down-regulated DEPs in Mp*fer-1* and Mp*mri-1* and the number of DEPs common to both analyses. The statistical significance of overlapping datasets in C) and D) was determined using a hypergeometric test.

Significantly enriched gene families (False Discovery Rate FDR <0.01) were identified using the GenFam online tool (https://www.mandadilab.com/genfam/ ; Bedre & Mandadi, 2019) using a Fisher’s exact test with Benjamini-Hochberg multiple testing correction (Figure 4A, B). Gene ontology (GO) enrichment analysis was also performed using the online tool PlantRegMap (http://plantregmap.gao-lab.org/go.php; Tian *et al*., 2020). Significantly enriched GO categories were selected using Fisher’s exact test with FDR <0.01 (Figure S3). For both GenFam and GO enrichment analyses, gene identifiers were converted from MpTakv6.1 to JGI3.1 prior to analysis using the ID converter tool from MarpolBase (https://marchantia.info/utils/id_converter/).

**Figure 4:**
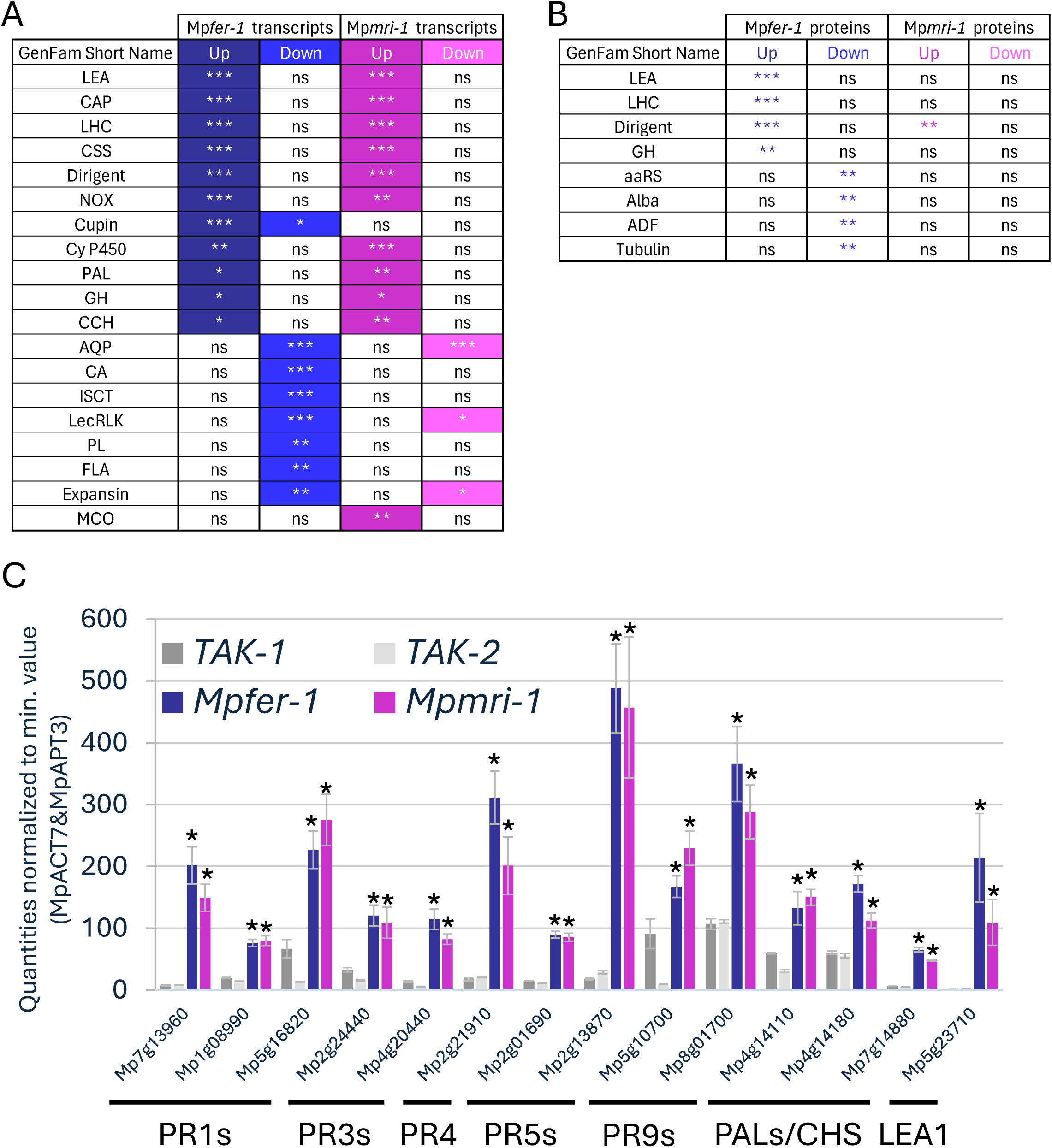
Gene family (GenFam) enrichment analysis and qRT-PCR validation of *PR*, *PAL* and *LEA* gene expression in the *Marchantia* CWI mutants. A) Table showing gene families significantly enriched in the transcriptomic data sets. B) Table showing gene families significantly enriched in the proteomic data sets. Three stars (***) indicates statistical significance p≤0.001, two stars (**) indicates p≤0.01, one star (*) indicates p≤0.05, and ‘ns’ indicates not significant. C) Quantitative real-time PCR results of 10 *PR* genes, 3 genes from the phenylalanine ammonia-lyase/chalcone synthase (*PAL/CHS*) gene family, and one gene from the late embryogenesis abundant (*LEA*) gene family. These genes and gene families were enriched in the transcriptome of both the Mp*fer-1* and Mp*mri-1* mutants. Normalization was performed against Mp*APT3* and Mp*ACT7*. (*) indicates significant difference between mutants and its WT strain at p≤0.1 according to the unified Wilcoxon–Mann–Whitney test.

### Quantitative Real-Time PCR (qRT-PCR)

One microgram of total RNA was reverse-transcribed (RevertAid First Strand cDNA Synthesis Kit, Thermo Fisher Scientific). qRT-PCRs were carried out in a QuantStudio 5 System (ABI/Life Technologies) using 96 well plates (0.2 ml) and cover foil (Opti-Seal Optical Disposable Adhesive) from BIOplastics. Reaction mixtures of 10 μl were composed of 5 μl GoTaq qRT-PCR Master Mix (Promega), 0.2 μl of each primer (10 μM), 1 μl diluted cDNA, and 3.6 μl ddH_2_O. Amplification was carried out with the standard settings of the QuantStudio 5 System (50°C for 2 min, 95°C for 10 sec, 40 cycles at 95°C for 15 sec, and 60°C for 1 min, followed by 95°C for 15 sec and a final dissociation curve from 60°C to 95°C). Analysis was performed using QuantStudio TM Design, Analysis Software version 1.4.1 and Excel 2019.

Sequences for primer design were taken from MarpolBase (Genome Database for *Marchantia polymorpha*, https://marchantia.info/), and *in silico* sequence analysis was carried out with CLC DNA Workbench version 5.6.1. The primers were designed using GenScript Real-time PCR Primer Design (www.genscript.com) at an optimum Tm of 60±2°C. All amplicons showed a single band of the expected size in gel electrophoresis and a single peak in the melting curve. Average Cq values (y) and log10 values of three biological and three technical replicates of six serial dilutions of 1:5, 1:10, 1:20, 1:40, 1:80, and 1:160 (x) were used to calculate the slope of a regression line using the formula 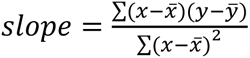. Efficiency of primers was then calculated with *efficiency*[%] = 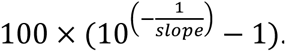. Correlation of the Cq values (R) was calculated with the formula 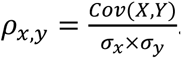. Primers for the reference genes Mp*APT3* and Mp*ACT7* were described before (Saint-Marcoux et al., 2015). Novel primers for genes of interest were accepted with an efficiency of 80–120% and a correlation between -1 and -0.99 (Table S1).

All transcript quantities of the genes of interest represent the average of four biological and three technical replicates from qRT-PCRs of 1:5 diluted cDNA samples. Outliers in the raw data were removed using a two-sided Grubbs test at a significance level of 0.05. Normalization against the two reference genes Mp*APT3* and Mp*ACT7* was carried out using normalization factors, according to the geNorm manual (Vandesompele *et al*., 2002).

Since exponentiation causes the data to be strictly positive, the standard normal assumption is not appropriate (Kirkwood, 1979). However, using a Kolmogorov-Smirnoff test on the log-transformed data, we were able to demonstrate that the data is log-Normally distributed. We use *1 + efficiency* as the base for our logarithm, leading to the adjusted formula 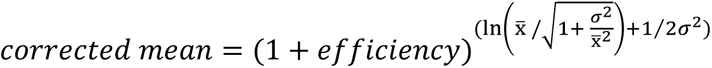 with σ^2^ = ln (1 + 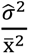) (variance). The bias-corrected standard deviations are given by *SD*[*X*] = *E*[*X*] ∗ *CV*[*X*] with 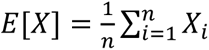 (empirical mean), 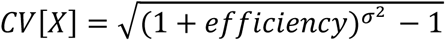 (coefficient of variation), and 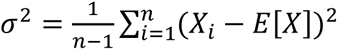 (empirical standard deviation).

### Proteomic Analyses

For protein extraction and fractionation, 4 biological replicates of frozen thalli with their rhizoids (3 thalli per replicate for Tak-1 and Tak-2 controls, 5-6 thalli for Mp*fer-1* and Mp*mri-1* smaller mutant plants), were disrupted in a Retsch mill with stainless steel beads for 5 min at 30 Hz. To the finely ground plant material 300 µL urea/DTT extraction buffer (8M urea/5mM DTT in 100mM Tris, pH 8.5/ protease inhibitor cocktail 2 and 3, Sigma, 20 µL/mL) was added, samples were mixed and incubated for 30 min at room temperature (RT). After centrifugation for 10 min (15k x g) at RT the supernatant was removed and transferred to a fresh tube, this procedure was repeated up to 3 times, until no visible pellet was retained. Protein concentration was determined using the Pierce 660 nm protein assay.

An aliquot of 50 µL total protein extract was subjected to a chloroform-methanol precipitation (Wessel & Flügge, 1984). In brief, 200 µL methanol were added, and samples were vortexed. Next, 50 µL chloroform were added and the samples were vortexed again. Finally, 150 µL H_2_O were added and, after vortexing, samples were centrifuged for 1 min at 14k x g. The upper layer was then removed, without disturbing the layer of precipitated protein. 200 µL of methanol were added, and the samples were vortexed and centrifuged for 10 min at 18k x g to pellet the precipitate. After removal of the organic layer the pellet was air-dried for 10 min and subjected to an in-solution digest.

Samples were taken up in 25 µL 8M urea/5 mM DTT (pH 8.5) and incubated for 30 min after which the samples were alkylated using CAA (550 mM, 0.6 µL), after 30 min samples were diluted with 175 µL 100mM Tris, pH 8.5, 1mM CaCl_2_, and 0.5 µg trypsin were added. Samples were incubated overnight shaking at 32°C. Samples were acidified with 5 µL 50% TFA and split into single BoxCar and library samples. For the library samples, 20 µL (5 µg) of each condition per replicate were mixed and submitted to SCX fractionation. To this end, StageTips were prepared using 6 layers of SPE disk (Empore Cation 2251 material) activated with acetonitrile and 1% TFA (100 µL each), washed with buffer A (water, 0.2% TFA, 100 µL) by spinning 5 min (1.5k x g), loaded by centrifugation (10 min, 800 x g), and washed with buffer A (5 min, 1.5k x g, 100 µL). Fractionation was carried out using an ammonium acetate gradient (20% ACN, 0.5 % FA) starting from 25 mM to 500 mM for 9 fractions and two final elution steps using 1% ammonium hydroxide, 80% ACN and finally 5% ammonium hydroxide 80% ACN. All fractions were eluted by centrifugation (5 min, 500 x g) using 2 x 30 µL eluent. The fractions were dried and taken up in 10 µL A* (2% ACN, 0.1% TFA) buffer. The peptide concentration was determined by Nanodrop and samples were diluted to 0.1 µg/µL for measurement. The first fraction was discarded, and samples were measured in reverse order of elution.

For the BoxCar analysis, the remaining samples were desalted with C18 Empore disk membranes according to the StageTip protocol (Rappsilber *et al*., 2003). The eluted peptides were dried and then taken up in 10 µL A* buffer. The peptide concentration was determined by Nanodrop, and samples were diluted to 1:10 for measurement.

For LC-MS/MS data acquisition, library samples were analyzed using an EASY-nLC 1200 (Thermo Fisher) coupled to a Q Exactive Plus mass spectrometer (Thermo Fisher). Peptides were separated on 16 cm frit-less silica emitters (Nikkyo Technos Co., Ltd., Japan, 75 µm inner diameter), packed in-house with reversed-phase ReproSil-Pur C18 AQ 1.9 µm resin (Dr. Maisch). Peptides were loaded on the column and eluted for 115 min using a segmented linear gradient of 5% to 95% solvent B (0 min : 5%B; 0-5 min -> 5%B; 5-65 min -> 20%B; 65-90 min ->35%B; 90-100 min -> 55%; 100-105 min ->95%, 105-115 min ->95%; solvent A 0% ACN, 0.1% FA; solvent B 80% ACN, 0.1%FA) at a flow rate of 300 nL/min. Mass spectra were acquired in data-dependent acquisition mode with a TOP15 method. MS spectra were acquired in the Orbitrap analyzer with a mass range of 300–1750 m/z at a resolution of 70,000 FWHM and a target value of 3×10^6^ ions. Precursors were selected with an isolation window of 1.3 m/z. HCD fragmentation was performed at a normalized collision energy of 25. MS/MS spectra were acquired with a target value of 10^5^ ions at a resolution of 17,500 FWHM, a maximum injection time (max.) of 55 ms, and a fixed first mass of m/z 100. Peptides with a charge of +1, greater than 6, or with an unassigned charge state were excluded from fragmentation for MS2. Dynamic exclusion for 30s prevented repeated selection of precursors.

BoxCar samples were analyzed using an EASY-nLC 1200 (Thermo Fisher) coupled to a Q Exactive Plus mass spectrometer (Thermo Fisher). Peptides were separated on 16 cm frit-less silica emitters (Nikkyo Technos Co., Ltd., Japan, 75 µm inner diameter), packed in-house with reversed-phase ReproSil-Pur C18 AQ 1.9 µm resin (Dr. Maisch). Peptides were loaded on the column and eluted for 115 min using a segmented linear gradient of 5% to 95% solvent B (0 min : 5%B; 0-5 min -> 5%B; 5-65 min -> 20%B; 65-90 min - >35%B; 90-100 min -> 55%; 100-105 min ->95%, 105-115 min ->95%; solvent A 0% ACN, 0.1% FA; solvent B 80% ACN, 0.1%FA) at a flow rate of 300 nL/min. Mass spectra were acquired in a data-independent manner using the MaxQuant.Live application (Wichmann *et al*., 2019). The acquisition was initiated using the “magic scan” protocol consisting of one full MS scan with a mass range of 300–1650 m/z at a resolution of 140,000 FWHM, a target value of 3×10^6^ ions and a maximum injection time of 20 ms. This was followed by two BoxCar scans, each consisting of 10 boxes with 1 Da overlap and a scan range from 400-1200 m/z. The maximum injection time for a BoxCar scan was set to 250 ms, with a resolution of 140,000 FWHM, and a target value of 5×10^5^ ions. The 5 most abundant ions from each BoxCar scan were selected for HCD fragmentation at a normalized collision energy of 27. Precursors were selected with an isolation window of 1.4 m/z. MS/MS spectra were acquired with a target value of 10^5^ ions at a resolution of 17,500 FWHM, a maximum injection time (max.) of 28 ms and a fixed first mass of m/z 50. Peptides with a charge of +1, greater than 5 or with an unassigned charge state were excluded from fragmentation for MS2. Dynamic exclusion for 30s prevented repeated selection of precursors.

The MS proteomics data have been deposited to the ProteomeXchange Consortium via the PRIDE partner repository with the dataset identifier PXD058089 (Perez-Riverol *et al*., 2022). Raw data were processed using MaxQuant software (version 1.6.3.4, http://www.maxquant.org/) (Cox & Mann, 2008) with label-free quantification (LFQ) and iBAQ enabled (Tyanova *et al*., 2016). Library samples and BoxCar samples were grouped into separate parameter groups. In the group specific parameters, library samples were set to “Standard” type and BoxCar samples to “BoxCar” type. In the Misc. setting the Match type for library samples was set to “match from” and for BoxCar to “match from and to”.

MS/MS spectra were searched by the Andromeda search engine against a combined database containing the sequences from *Marchantia* Tak-1 and Tak-2 genotypes (MpTakv6.1) and sequences of 248 common contaminant proteins and decoy sequences. Trypsin specificity was required and a maximum of two missed cleavages were allowed. The minimal peptide length was set to seven amino acids. Carbamidomethylation of cysteine residues was set as fixed, oxidation of methionine and protein N-terminal acetylation were set as variable modifications. The match between runs option was enabled, and the match-time window was set to 1.5 min. Peptide-spectrum-matches and proteins were retained if they were below a false discovery rate (FDR) of 1% in both cases.

Statistical analysis of the MaxLFQ values was carried out using Perseus (version 1.5.8.5, http://www.maxquant.org/). Quantified proteins were filtered for reverse hits and hits “only identified by site” and MaxLFQ values were log2 transformed. After grouping samples by condition only those proteins with three valid values in one of the conditions were retained for the subsequent analysis. Two-sample t-tests were performed using a permutation-based FDR of 5%. Alternatively, quantified proteins were grouped by condition and only hits with 4 valid values in one of the conditions were retained. Missing values were imputed from a normal distribution, using the default settings in Perseus (1.8 downshift, separately for each column). Volcano plots were generated in Rstudio with the *ggplot2* package (Figure S1B and Figure 2B, 3B).

### ABA-related responses in *Marchantia*

For growth quantification, images of at least twenty individual thalli were collected using a Scanner Epson Perfection V850 Pro. Whole thallus growth was quantified by measuring the area of plants grown from gemmae over a period of 14 days. Thallus surface area was determined using classical segmentation by color-thresholding with the Fiji-ImageJ v1.46 program (Schindelin *et al*., 2012). The area growth rate (%) was calculated as a fraction of the average of ABA-treated plants’ surface area to the average of Mock-treated surface area. Graph design and statistical analyses were performed using the GraphPad Prism 8 software (GraphPad Prism Software Inc.). Statistically significant differences were determined by two-way analysis of variance (ANOVA), followed by Tukey’s method with a significance level of p ≤0.05.

Concerning ABA-related qRT-PCR, total RNA was prepared from plant tissues using Promega’s SV Total RNA Isolation System according to the supplier’s instructions including an RNase-Free DNase I treatment. The SV Total RNA Isolation System produced typical yields of approximately 50-400ng/µL. A total of 400 ng of RNA was reverse transcribed using SuperScript IV reverse transcriptase (Thermo Fisher Scientific) according to the supplier’s instructions. The cDNAs were diluted 10-fold, and 1.5 μl of this dilution was added to 5 μl of qRT-PCR Brilliant III SYBR Master Mix (Agilent Technologies). Quantitative Real-time PCR (qRT-PCR) was performed on an AriaMx Real-time PCR System (Agilent Technologies) with a program consisting of 3 min of initial denaturation at 95°C, followed by 40 cycles consisting of 5 s at 95°C and 10 s at 60°C. Primers used for PCR experiments are listed in Table S1. All data were from three independent biological replicates each involving three technical replicates. Reference genes were Mp*ACT* and Mp*APT3*, as indicated in the figure legends. Data from AriaMx v1. 0.8 were quantified with the comparative delta-Ct method, using the individual primer efficiency corrected calculation (Rao *et al*., 2013). Graph design and statistical analyses were performed using the GraphPad Prism 8 software (GraphPad Prism Software Inc.). Statistically significant differences for qRT-PCR data were determined by two-way analysis of variance (ANOVA), followed by Tukey’s method with a significance level of p ≤0.05.

## RESULTS

### The transcriptome and proteome of cell wall integrity-related mutants Mp*fer-1* and Mp*mri-1* overlap significantly

To reveal pathways that could be regulated by the ancestral FER/MRI module or alternative compensatory ones that are de-regulated in the absence of the module, comparative transcriptomic (RNA-seq) and proteomic (LC-MS/MS) analyses of Mp*fer-1* (male) and Mp*mri-1* (female) lines and their respective wild-type male (Tak-1) and female (Tak-2) strains were carried out. For this purpose, 3-week-old *Marchantia* thalli grown on cellophane membranes were collected and RNA and proteins were extracted (Figure 1A).

As seen in Figure 1, principal component analyses (PCAs) showed robust clustering indicating reproducibility between our independent biological samples for the transcriptomes (n=3, Figure 1B) and the proteomes (n=4, Figure 1C), as well as a clear separation between both mutants and the wild-type strains.

While the PCA indicated some similarity between Tak-1 and Tak-2 transcriptomes, we decided to compare the female Tak-2 to the male Tak-1 thalli transcriptomes. As expected, this analysis identified mostly, but not exclusively, transcripts of genes located on the female U and male V chromosomes among the top differentially expressed genes (Fig. S1, see how U and V chromosomes located genes are on both ends of the volcano plot). First, this confirms that sex chromosomes located genes are also expressed in vegetative tissues, maybe as relics of the ancestral autosome from which the sex chromosomes evolved (Bowman *et al*., 2017). Secondly, it indicates that a subset of genes located on autosomes are under the transcriptional control of sex chromosome-encoded genes even in vegetative tissues. For instance, this is the case for the feminizer gene *BASIC PENTACYSTEINE ON THE U* (*BPCU*, MpUg00370, 24^th^ most up-regulated gene in Tak-2 vs Tak-1), which regulates the autosomal transcription factor *FEMALE GAMETOPHYTE MYB* (*FGMYB*, Mp1g17210, 40^th^ most down-regulated gene of Tak-2 vs Tak-1), which in turn promotes female sexual differentiation (Iwasaki *et al*., 2021).

Concerning the cell wall integrity mutants, transcriptome profiling identified 2,293 and 2,038 transcripts that are differentially expressed (log2 (FC)≥1.5; p-value ≤ 0.05) in Mp*mri-1* vs Tak-2 and Mp*fer-1* vs Tak-1, respectively (Figure 2). Out of roughly 5,200 quantified proteins, our proteomic analysis determined that 775 and 1,549 proteins have different abundance (log2 (FC)>1.5; p-value < 0.05) in Mp*mri-1* vs Tak-2 and Mp*fer-1* vs Tak-1, respectively (Figure 3). For both mutants, there were more up-regulated than down-regulated transcripts (1,572 and 1,127 up-vs 721 and 911 down-regulated in Mp*mri-1* and Mp*fer-1*, respectively), and more enriched proteins than depleted ones (526 and 955 entiched vs 249 and 594 depleted in Mp*mri-1* and Mp*fer-1*, respectively). This indicates that, globally, the MpFER/MpMRI module negatively regulates transcriptional/translational networks.

The transcriptome and proteome data of Mp*fer-1* significantly overlapped (Figure S2A, B), which was true for Mp*mri-1* as well, although to a lesser extent (Figure S2C, D). More interestingly, large and significant overlaps were observed between DEGs in Mp*fer-1* and Mp*mri-1* (Figure 2 C, D), as well as between differentially abundant peptides in Mp*fer-1* and Mp*mri-1* (Figure 3C, D) highlighting that MpFER and MpMRI cooperation is not solely limited to the maintenance of CWI in rhizoids (Westermann *et al*., 2019).

### Transcriptome and proteome comparisons indicate that the MpFER/MpMRI module regulates large gene families related to ABA and defense responses

We performed gene ontology (GO) analysis on both the differentially expressed genes (DEGs) and proteins (DEPs) using PlantRegMap (https://plantregmap.gao-lab.org/go.php). It revealed some overlap in categories found in the transcriptomic and proteomic data sets (Figure S3), including the GO terms chitin binding (GO:0008061), catalytic activity (GO:0003824), and hydrolase activity - both hydrolysing O-glycosyl compounds (GO:0004553) and acting on glycosyl bonds (GO:0016798). However, most of the GO terms identified by this analysis were too vague to help us infer precise functions for MpFER or MpMRI. More importantly, when we compared our list of GO terms with those from similar studies evaluating AtFER (Zhao *et al*., 2021; Wang *et al*., 2022) we found very few similarities suggesting that AtFER may have acquired additional functions in the angiosperm lineage or that the available GO database for Marchantia is not yet as comprehensive as for Arabidopsis. However, the presence of terms related to CW (GO:0005618, GO:0005576, GO:0071944, GO:0048046) and responses to oxidative stress (GO:0006979) suggest MpFER and MpMRI share some common functionality with their Arabidopsis homologs.

Next, we performed GenFam enrichment analysis on all DEGs and DEPs identified in the CWI mutant vs WT strains (Figure 4A, B). Families of aquaporins, lectin-RLKs and expansins were significantly enriched in down-regulated genes of both mutants while those of LEA (Late Embryogenesis Abundant, ABA-responsive genes), CAP (Cysteine rich secretory proteins, e.g., Pathogenesis Related-1, PR1), LHC (Light-Harvesting Chlorophyll binding), CSS (Chalcone and Stilbene Synthesis), Dirigent (DIR; derived from the Latin word, dirigere, *to align or guide*), PAL (Phenylalanine Ammonia Lyases, enzymes that eliminate ammonia from phenylalanine to form cinnamate, a precursor of lignins/lignans flavonoids, and thus anthocyanins), NOX (NADPH oxidase), Cytochromes P450, GH (Glycosyl Hydrolase), and CCH (Copper Chaperone) were enriched in up-regulated genes of both mutants (Figure 4A). Interestingly, LEA, LHC, DIR and GH were also enriched in more abundant proteins of Mp*fer-1* (Figure 4B), while DIR was enriched in up-regulated transcripts and more abundant proteins of both Mp*fer-1* and Mp*mri-1*. GHs are carbohydrate-active enzymes (CAZymes) that can selectively cleave various carbohydrates and glycoconjugates with functions such as CW remodelling, reserve mobilization and pathogen defense (Kfoury *et al*., 2024). DIR proteins are thought to control the stereoselective coupling of coniferyl alcohol in the formation of lignan and lignin (Davin *et al*., 1997). Many DIR encoding genes are inducible by different types of abiotic and biotic stress factors, including wounding, dehydration, low temperature, ABA, H_2_O_2_, NaCl (Paniagua *et al*., 2017) or following Colletotrichum, Fusarium and Phytophthora infection (Ponce De León *et al*., 2012; Reboledo *et al*., 2015; Carella *et al*., 2019). The predominant roles of the *DIR* gene family are in secondary metabolism and defense responses and recently some members were shown to regulate the localized polymerization of lignin required for the formation of the Casparian strip in the cell wall and for attaching the strip to the plasma membrane in Arabidopsis (Gao *et al*., 2023a). In *Marchantia*, approximately half of the DIR were found in the cell wall proteome (Kolkas *et al*., 2021), suggesting that the CW of Mp*fer-1* and Mp*mri-1* are likely quite different from the ones of wild-type plants. LEAs are a large and diverse group of small hydrophilic disordered proteins involved in protecting plant tissues and cells from abiotic stresses and low water availability. They are present from algae to angiosperms and their expression levels are rapidly and highly up-regulated by salt, cold, heat, drought and ABA treatments (Hsiao, 2024).

Based on the GenFam enrichment analysis, we therefore hypothesized that gene families for PRs, DIRs, CAPs, PALs, Chitinases, Anthocyanin biosynthesis as well as ABA-related genes may globally be mis-regulated in the *Marchantia* CWI mutants. Indeed, when taking a closer look in our RNA-seq data at the expression levels of these toolbox genes across all our biological replicates, they did show globally higher expression levels in Mp*fer-1* and Mp*mri-1* mutant samples as compared to wild-type ones, thereby supporting the GenFam analysis (Figure S4-S7). Moreover, this was independently confirmed by picking representative genes for qRT-PCR analyses which showed significantly higher expression levels for *PR1, 3, 4 5, 9*, *PAL/CHS* and *LEA1* in the mutants compared to wild-type plants (Figure 4C).

Altogether, our results indicate that MpFER and MpMRI globally exert a negative regulation on defense-responsive genes and ABA responses as observed for Arabidopsis AtFER (Wang *et al*., 2022).

### Mp*fer-1* but not Mp*mri-1* plants exhibit priming and hypersensitivity to ABA exposure

Since our multi-omics data pointed to a role for MpFER and MpMRI in regulating ABA responses (Figure 4 and S7B), we compared our transcriptomes of Mp*fer-1* and Mp*mri-1* with that of Tak-1 plants treated with 1 μM ABA for 6 h (Jahan *et al*., 2019). Interestingly, while there was little overlap in down-regulated genes (Figure S8A, B), there was a highly significant overlap between up-regulated genes in Mp*fer-1*/Mp*mri-1* vs up-regulated genes in ABA-treated Tak-1 (Figure S8C, D). This indicates that a significant pool of genes that are up-regulated by ABA treatment is globally already up-regulated in CWI-related mutant plants without ABA treatment, as confirmed for Mp*LEA1* (Figure 4C) and other ABA-related genes (Figure S7B).

As At*fer-4* plants were shown to be hypersensitive to ABA in seed germination, stomatal and root growth assays (Yu *et al*., 2012; Chen *et al*., 2016), we next tested Mp*fer-1* and Mp*mri-1* mutant plant responses to a 14 day-treatment of 0, 1 and 10 μM ABA. Analyses of plant growth responses show that surprisingly, among the 4 genotypes, Tak-1 was the least affected. Indeed, for all genotypes but Tak-1, ABA-induced growth inhibition was already significant at 1 μM ABA and became more pronounced at 10 μM ABA (Figure 5 and S9 for an independent assay). Tak-1 plants only showed a mild response at 10 μM ABA. In clear contrast, Mp*fer-1* growth inhibition was the highest among all genotypes at both 1 and 10 μM ABA (Figure 5 and S9). These results indicate that Mp*fer-1* plants are hypersensitive to ABA treatment while Mp*mri-1* behaves like Tak-2.

**Figure 5:**
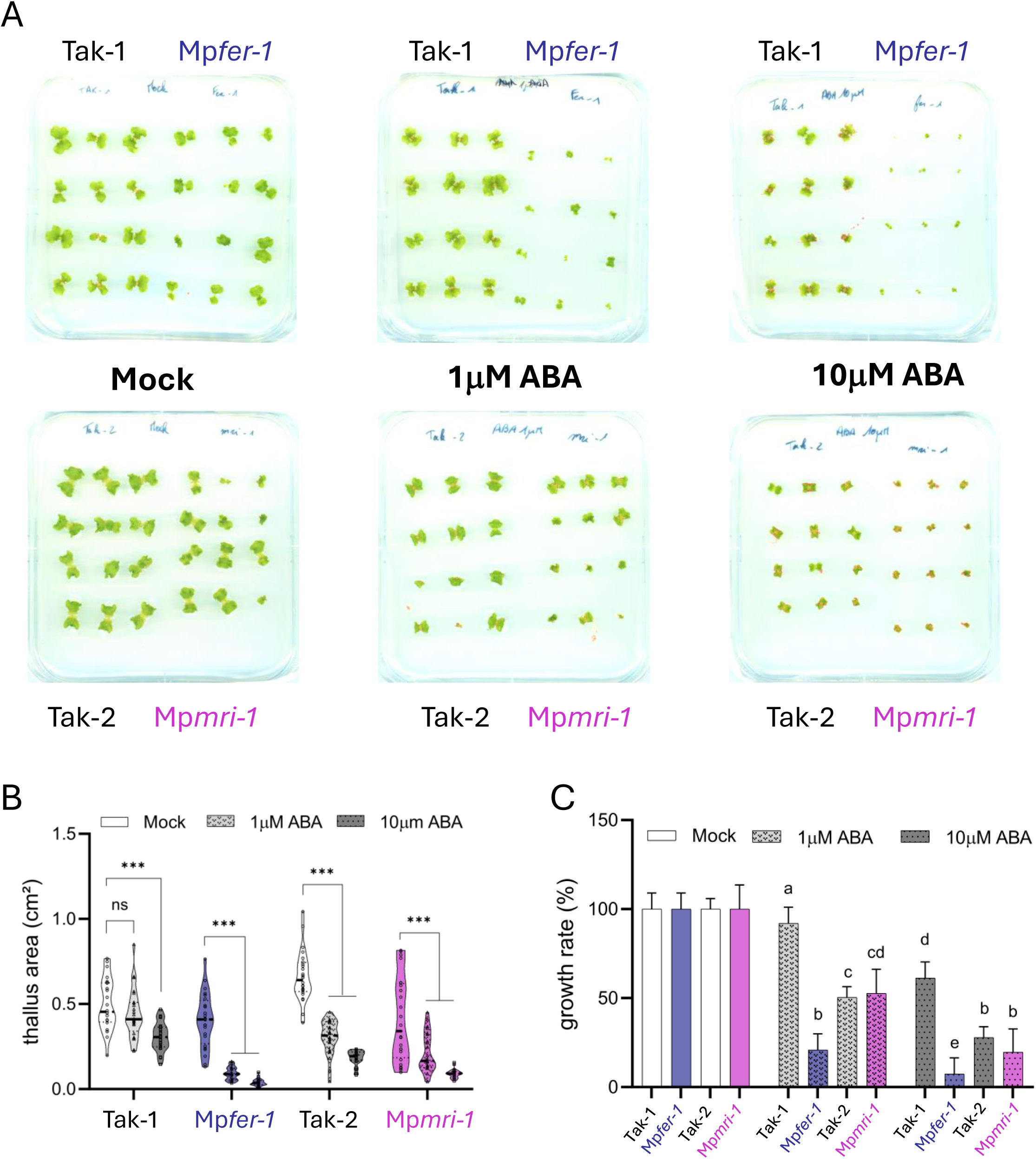
ABA treated *Marchantia* CWI mutants and quantification of thallus area and growth rate under ABA treatment. A) Representative images of *Marchantia* Tak-1, Tak-2, Mp*fer-1*, and Mp*mri-1* genotypes grown on agar plates supplemented with 0 μM ABA (mock), 1 μM ABA, or 10 μM ABA. B) Violin plot showing the thallus area of the four genotypes grown with 0 μM ABA (mock), 1 μM ABA, or 10 μM ABA. C) Bar chart showing the calculated growth rate as a fraction of the average of ABA-treated plant surface area to the average of Mock-treated surface area. Statistically significant differences were determined by two-way analysis of variance (ANOVA), followed by Tukey’s method with a significant level of P ≤0.05. See also Figure S9 for an independent assay.

To further understand the link between MpFER and ABA, we quantified by qRT-PCR transcript levels for ABA-related genes including ABA biosynthesis (Mp*ABA1*, Mp*NCED*), catabolism (Mp*CYP707A*), receptor and early signalling (Mp*PYL1*, Mp*SnRK2A*), transcription factors (Mp*ABI3A*, Mp*ABI5A*), responsive genes (Mp*LEA1L*, Mp*Dehyhdrin*) and regulator (Mp*MFT*) in thalli grown 14 days on 10 μM ABA. The Type 2C protein phosphatase Mp*ABI1A* (MpVg00730), the main negative regulator of ABA signalling was not tested as it is located on the male chromosome and its transcripts would not have been detected in the female lines. In addition, to Tak-1, Mpf*er-1*, Tak-2 and Mp*mri-1*, we included 2 additional lines JW26-3 and JW26-13. These lines are progenies segregating from a Mp*fer-1* (male) x Tak-2 (female) cross and are both female with JW26-3 not harbouring the T-DNA insertion of Mp*fer-1* while JW26-13 still carries it. In agreement, JW26-3 plants displayed WT-like thalli sizes and carried intact rhizoids while JW26-13 plants were much smaller with burst rhizoids as for Mp*fer-1* (Figure S10).

Apart from Mp*CYP707A*, whose levels were moderately upregulated in Mp*mri-1* compared to wild-type strains (Figure 6A and Figure S7B), the T-DNA insertion in Mp*MRI* did not appear to influence ABA-related transcripts levels during this 14-day treatment with 10 μM ABA (Figure 6 and S11). This is in agreement with the normal responses to ABA treatment observed in Mp*mri-1* (Figure 5) and the globally lower expression levels of the ABA toolbox in Mp*mri-1* samples as compared to Mp*fer-1* ones (Figure S7).

**Figure 6:**
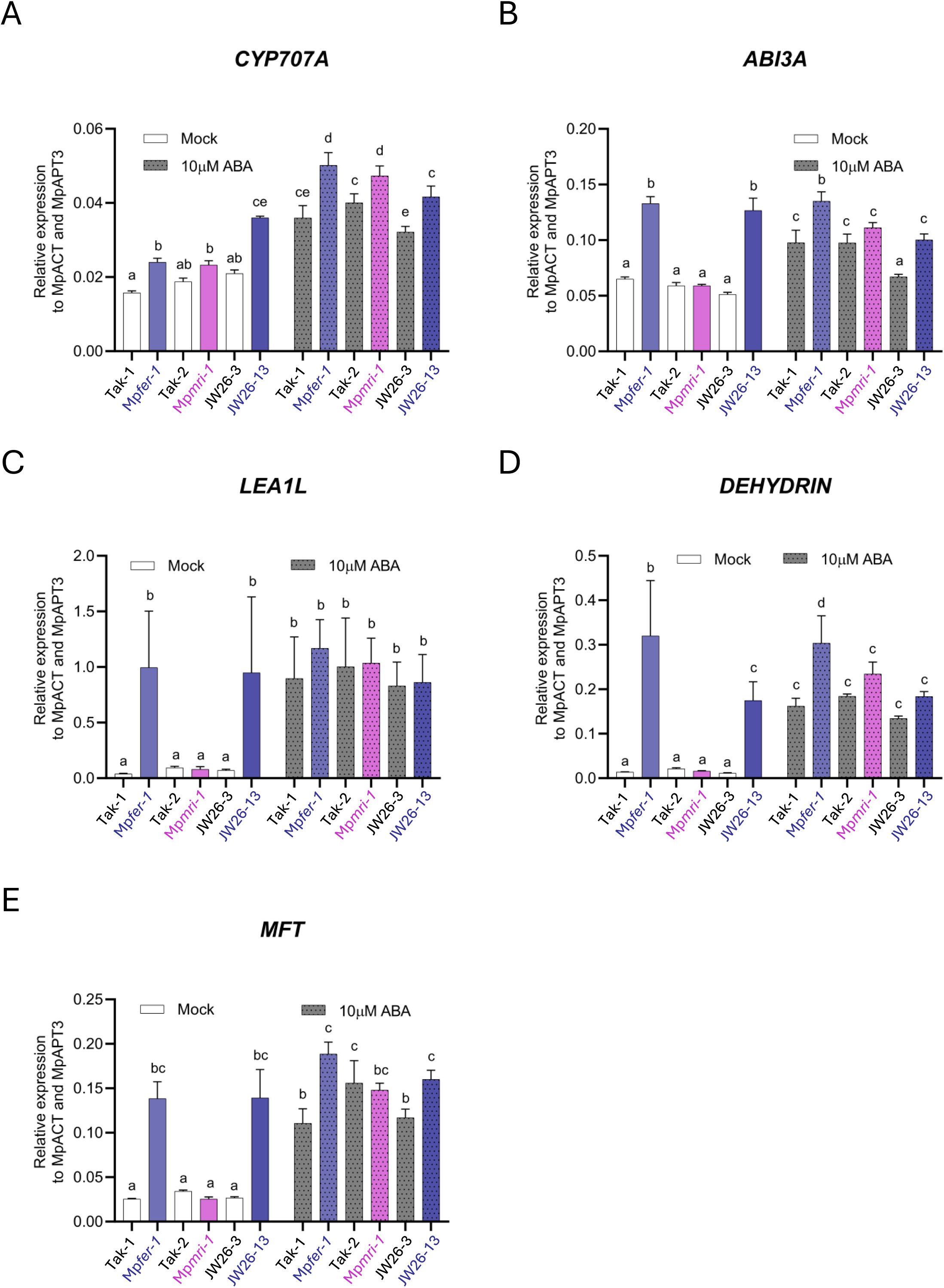
Gene expression analysis in Tak-1, Tak-2, Mp*fer-1,* Mp*mri-1,* JW26-3, and JW26-13 genotypes grown for 14 days on 10 μM ABA using quantitative real-time PCR (qRT-PCR). Transcripts levels for ABA-related genes were quantified, including A) the ABA catabolism gene Mp*CYP707A*, B) the ABA transcription factor Mp*ABI3A*, C) the ABA responsive genes Mp*LEA1L* and D) Mp*Dehyhdrin*, and E) the ABA regulator Mp*MFT*. Normalization was performed against Mp*APT3* and Mp*ACT7*. Statistically significant differences were determined by two-way analysis of variance (ANOVA), followed by Tukey’s method with a significant level of P ≤0.05.

Concerning Mp*fer-1*, firstly and as expected, across all experiments, the *fer-1* female mutant progeny JW26-13, behaved very similarly to the original male Mp*fer-1* (Figure 6 and S11), indicating that the sexual chromosomes do not influence the Mp*FER*-dependent ABA responses. Secondly, for Mp*fer-1* and its progeny JW26-13, ABA-related genes could be grouped into three categories. The first one includes genes whose levels were not impacted by the genotype of our mutant lines (e.g. Mp*ABA1* and Mp*ABI5*; Figure S11A, E). The second one includes genes that are slightly up-regulated when Mp*FER* is disrupted in mock conditions and do not increase upon the 14-days long ABA treatment (Mp*NCED*, Mp*PYL1*, Mp*SnRK2A*; Figure S11, B-D). The third one is the most prominent and includes genes that are up-regulated when Mp*FER* is disrupted (in Mp*fer-1,* male and JW26-13, female) in mock conditions and whose levels in all other genotypes are increased by the 14-days long ABA treatment to roughly the levels found in Mp*fer-1*/JW26-13 (Figure 6). This group comprises Mp*CYP707A*, Mp*ABI3A*, Mp*LEA1*, Mp*DHDN*, and Mp*MFT* and likely represents the targets of Mp*FER*-dependent inhibition. Their higher transcript levels in unchallenged Mp*fer-1* plants likely reflect an induced ABA state, explaining why Mp*fer-1* plants respond more strongly to even low exogenous ABA concentrations (Figure 5).

## DISCUSSION

In this study, comparative transcriptomic (RNA-seq) and proteomic (LC-MS/MS) analyses of CWI mutant lines vs wild-type strains showed that MpFER and MpMRI are global negative regulators of transcription and translation networks with roughly 50% more up-regulated transcripts and 66% more abundant proteins than down-regulated transcripts and depleted proteins in the mutants, respectively (Figures 2 and 3). Furthermore, our study reveals that more than half of the differentially expressed transcripts and abundant proteins identified in Mp*mri-1* mutant plants are also differentially expressed/abundant in Mp*fer-1* plants indicating that MpFER and MpMRI largely cooperate, but also most likely play specific roles independently of each other.

MpFER and MpMRI have been shown to genetically interact to maintain cell wall integrity during rhizoid growth (Westermann *et al*., 2019; Mecchia *et al*., 2022) just as it was shown for AtFER and AtMRI during root hair growth and AtANXUR1/2 and AtMRI during pollen tube growth (Boisson-Dernier *et al*., 2015). In Arabidopsis, AtFER’s role is far from being limited to tip-growing cells as it is crucial for almost all aspects of plant biology and growth in an ever-changing environment including ABA and defense responses (e.g. (Franck *et al*., 2018; Cheung, 2024)). In contrast, AtMRI has only been shown to be involved in regulating root hair and pollen tube growth so far (Boisson-Dernier *et al*., 2015; Liao *et al*., 2016). AtMRI belongs to the RLCK-VIII (also known as CARK for Cytosolic ABA Receptor Kinase) subfamily with 11 members in Arabidopsis, which have been reported to form heterodimers and homodimers with each other in planta (Li *et al*., 2022). On the one hand, MAZZA (CARK5) was shown to interact with CLAVATA1-like receptors and appeared to play a positive role in CLE peptide signalling in roots and possibly in stomata spacing (Blümke *et al*., 2021). On the other hand, CARK1, 3 and 6 have been shown to function as positive regulators of ABA-dependent physiological responses (Zhang *et al*., 2018; Wang *et al*., 2019, 2020) phosphorylating ABA receptors and inducing their de-dimerization (Li *et al*., 2022). Interestingly, AtMRI (CARK4) can also interact with and phosphorylate ABA receptors (Li *et al*., 2022), but no ABA-related phenotype has yet been described for AtMRI-related mutants. This could mean that either AtMRI plays no role during ABA signaling or that its ABA-related function is hidden by other closely related CARKs and the right multiple *cark* mutant combination has not been yet investigated. In *Marchantia*, the role of MpMRI appears to be more specific to governing rhizoid growth (Westermann *et al*., 2019), while a broader involvement of MpFER at both vegetative and reproductive stages of development was documented (Mecchia *et al*., 2022).

The large overlap between differentially expressed transcripts and abundant proteins observed in Mp*fer-1* and Mp*mri-1* displays a significant enrichment for gene families involved in responses to biotic and abiotic stresses in particular the *LEA*, *DIR*, *PR*, *GH*, *PAL*, *CHS* and *CSS* gene families (Figure 4A). Those enrichments can be visualized by looking at the heatmap of specific gene toolboxes across our samples (*e.g., PRs/PAL/CHS* Figure S5B), which were further confirmed by qRT-PCR analyses (Figure 4B).

Several gene families found to be up-regulated in the CWI mutants are related to plant defense responses indicating either that the MpFER/MpMRI module negatively regulates immunity or that the disruption of CWI and its impact on the CW physico-chemical properties activate autoimmune responses. In flowering plants such as Arabidopsis, soybean, rice, and tomato, FER has been shown to play contrasting roles in responses to pathogens. On the one hand, it was found to be a susceptibility factor for the obligate biotrophic fungi *Erysiphe orontii,* the agent of powdery mildew (Kessler *et al*., 2010), the hemibiotrophic fungi *Fusarium oxysporum* (Masachis *et al*., 2016; Fan *et al*., 2024) and *F. graminearum* (Wang *et al*., 2024), the hemibiotrophic fungi *Magnaporthe oryza*e (Yang *et al*., 2020) as well as for the parasitic nematode *Meloidogyne incognita* (Zhang *et al*., 2020, 2021). In the case of Fusarium and Meloidogyne, the resistance observed in the *fer*-related mutant was linked to RALF-like peptides secreted by the pathogens so as to manipulate FER-dependent pathways (Masachis *et al*., 2016; Zhang *et al*., 2020). On the other hand, FER has also been shown to play a positive role in immune responses to the bacterium *Pseudomonas syringae* acting at the plasma membrane as a scaffold component regulating the formation of immune receptor complex (Stegmann *et al*., 2017) and its mobility (Gronnier *et al*., 2022) and in the cytosol by destabilizing MYC2, a major positive regulator of jasmonic acid and coronatine responses (Guo *et al*., 2018). Excitingly, Chen *et al*. recently discovered that Arabidopsis infection by *P. syringae* or RALF23 treatment leads to cleavage of the cytoplasmic domain of FER and its translocation to the nucleus thereby likely promoting immune responses (Chen *et al*., 2024). In bryophytes, the role of MpFER during plant-pathogen interactions has not yet been characterized, but natural fungal pathogens have been isolated for *Marchantia* (Matsui *et al*., 2020) and pathosystems have been established with the oomycete *Phytophthora palmivora* (Carella *et al*., 2019), the bacteria *P. syringae* (Gimenez-Ibanez *et al*., 2019; Matsumoto *et al*., 2022), the fungi *F. oxysporum* (Redkar *et al*., 2022) and the tobacco mosaic virus TMV (Ros-Moner *et al*., 2024). Therefore, it will be possible to investigate whether the ancestral role of FER was to be a susceptibility factor, a positive regulator of immune responses or both depending on the pathogens.

Besides plant defense responses, the drought stress hormone ABA is also a common denominator known to regulate the expression of the gene families found up-regulated in Mp*fer-1* and Mp*mri-1*. In agreement, around 80% of up-regulated transcripts in wild-type *Marchantia* thalli treated with 1 μM ABA (Jahan *et al*., 2019) were found up-regulated in untreated CWI-related mutants (Figure S8). This suggested that the MpFER/MpMRI module negatively regulates ABA signaling.

However, despite displaying increased transcript levels of Mp*LEA1* (Figure 4C) and Mp*CYP707A* (Figure 6A), Mp*mri-1* plants displayed wild-type-like responses to ABA for thalli growth inhibition assays (Figure 5 and Figure S9). This indicates that MpMRI is not involved in ABA-dependent thalli growth inhibition and that the mis-regulation of ABA-related genes in Mp*mri-1* may be linked to a different, yet to be revealed, ABA function. In contrast, Mp*fer-1* exhibited enhanced expression of ABA-related genes compared to wild-type plants, including those coding for ABA biosynthesis (Mp*NCED*), an ABA receptor (MpP*YL1*), key positive regulators of ABA signaling (Mp*SnRK2A*, Mp*ABI3*, Mp*MFT*), and ABA responsive genes Mp*LEA1*, Mp*LEA1L* and Mp*Dehyhdrin* (Figure 4C, 6 and Figure S11). Interestingly, the genes of this group are also found to be downregulated by ectopic expression of the A-type protein phosphatase 2C PP2C Mp*ABI1A* (Eklund *et al*., 2018) suggesting that, MpFER and PP2C MpABI1A function together to negatively regulate ABA signaling. Consistently, we found that Mp*fer-1* plants were hypersensitive to ABA in thalli growth inhibition assays (Figure 5 and Figure S9). In Arabidopsis, AtFER has also been shown to negatively inhibit ABA signaling with At*fer-4* plants displaying hypersensitivity to ABA in seed germination, cotyledon greening, stomatal and root growth assays (Yu *et al*., 2012; Chen *et al*., 2016; Wang *et al*., 2022). In root growth assays, AtRALF1 also negatively impacted ABA response (Chen *et al*., 2016). It was shown that AtFER interacts with and activates the PP2C AtABI2, a key ABA negative regulator. Conversely, AtABI2 was shown to dephosphorylate AtFER and negatively feedback on ABA-activation of AtFER (Chen *et al*., 2016). In agreement, integrated omics on At*fer-4* mutant plants showed that the GO term “response to ABA” (GO:0009737) was significantly enriched among transcripts and proteins with increased levels in At*fer-4* (Wang *et al*., 2022). Moreover, FER was reported to phosphorylate and destabilize ABI5, a transcription factor and positive regulator of ABA inhibition of cotyledon greening in seedlings (Wang *et al*., 2022).

Altogether, our findings indicate that FER’s function of negatively regulating ABA signaling, likely in cooperation with A-type PP2Cs, is conserved between bryophytes and flowering plants and probably was present in the common ancestor of land plants. Furthermore, there are 3 RALFs in Marchantia that belong to the AtRALF1-clade of the RALF Family and 2 LLGs that could also constitute MpFER coreceptors (Mecchia et al., 2022). Assessing whether they could also play a negative role in ABA signaling would provide fundamental insights into the molecular mechanisms linking ABA signaling and cell wall integrity.

## ACKNOWLEDGEMENTS

We thank the Cologne Center for Genomics (<u>Cologne Center for Genomics (CCG)</u>) for generating the RNA libraries and performing the RNA-seq of our samples. We are grateful for the invaluable technical help provided by Roswitha Lentz, Sabine Lohmer and Birgit Kernebeck at the University of Cologne, and exploratory experiments carried out by Alexandre Brunet and Mathieu Brisson at the Institute Sophia Agrobiotech. We would like to offer our sincere appreciation to the IPO team for its incredible support.

---

**Table S1:**
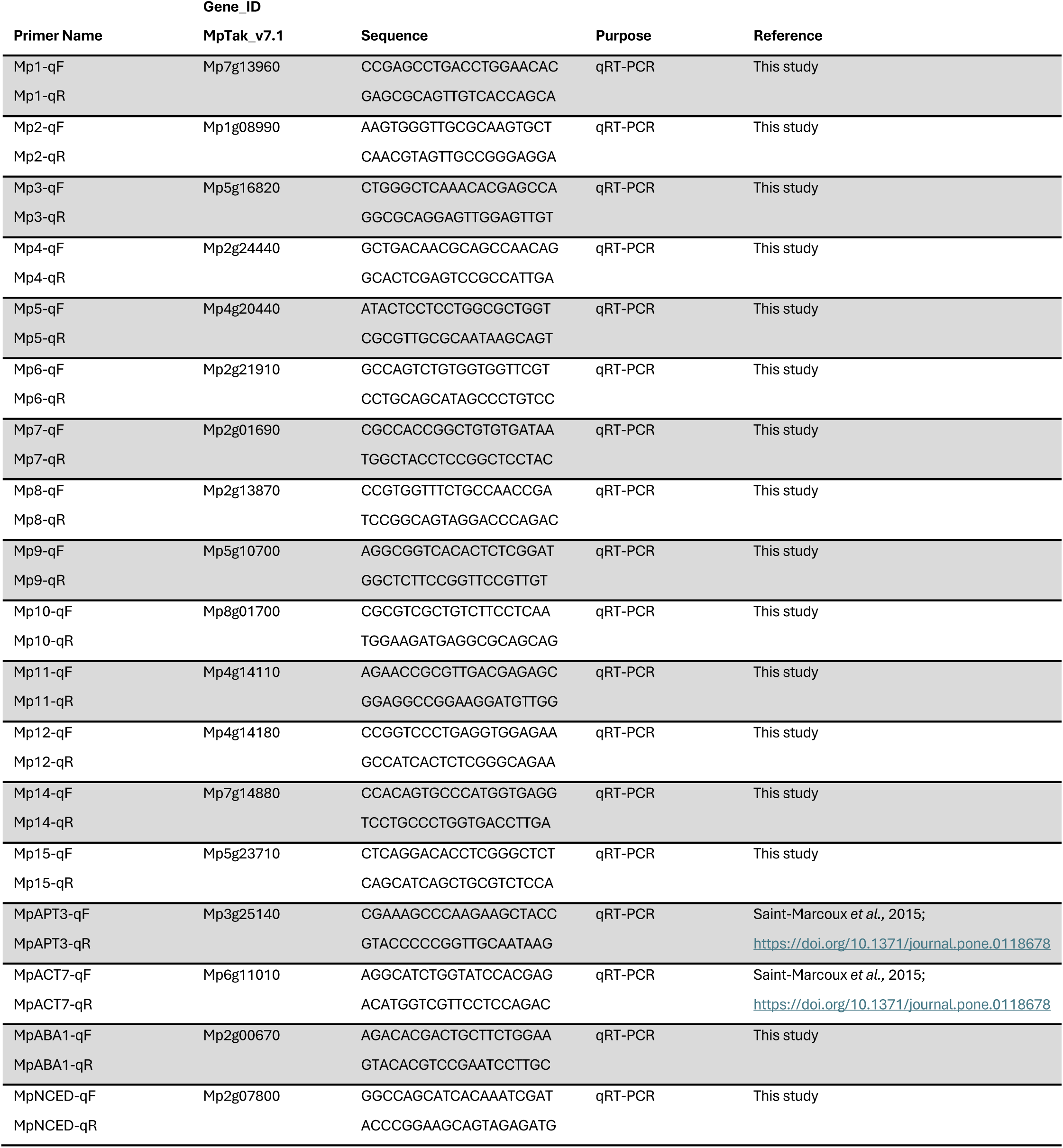
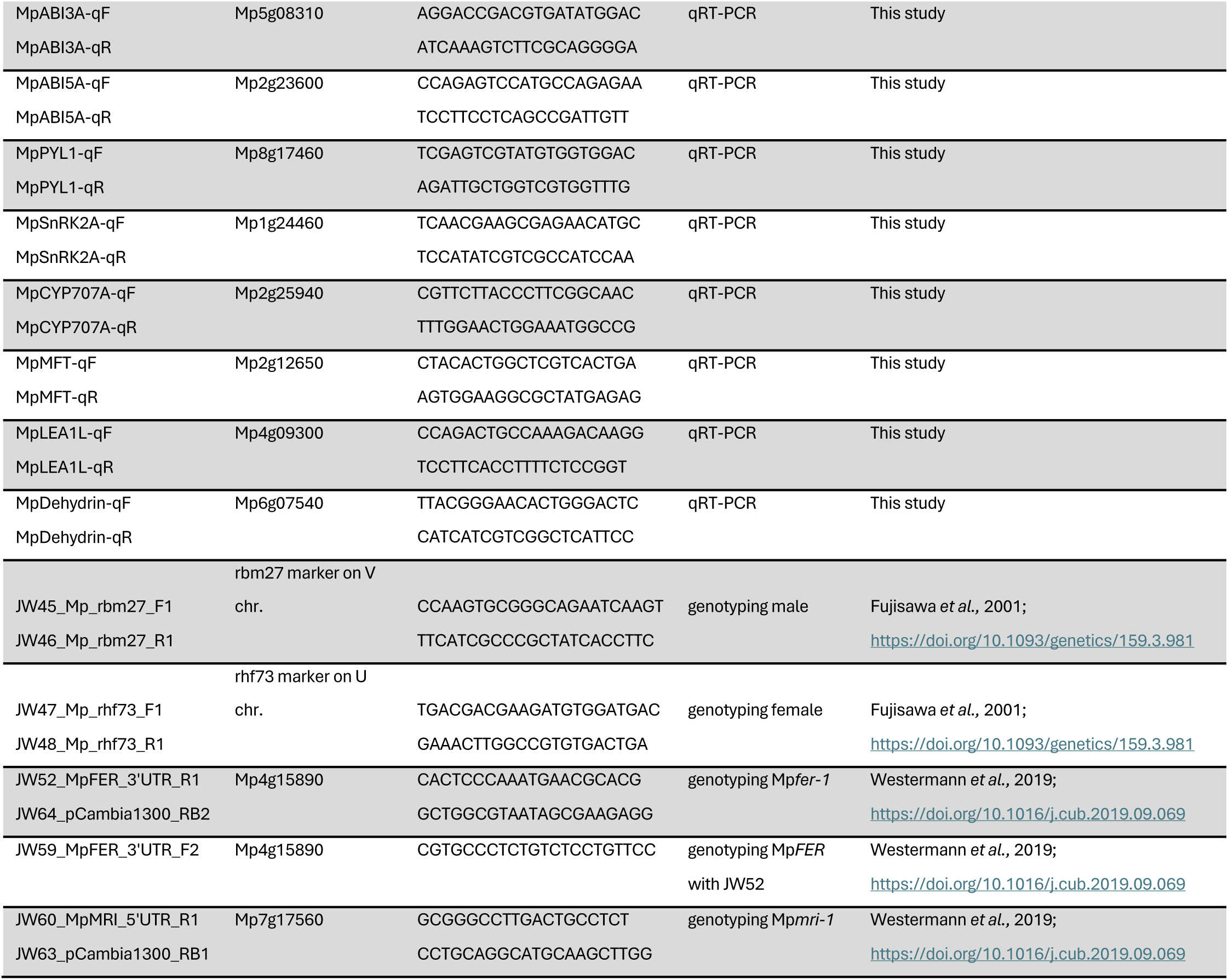
Oligonucleotides used in this study.

**Figure S1:**
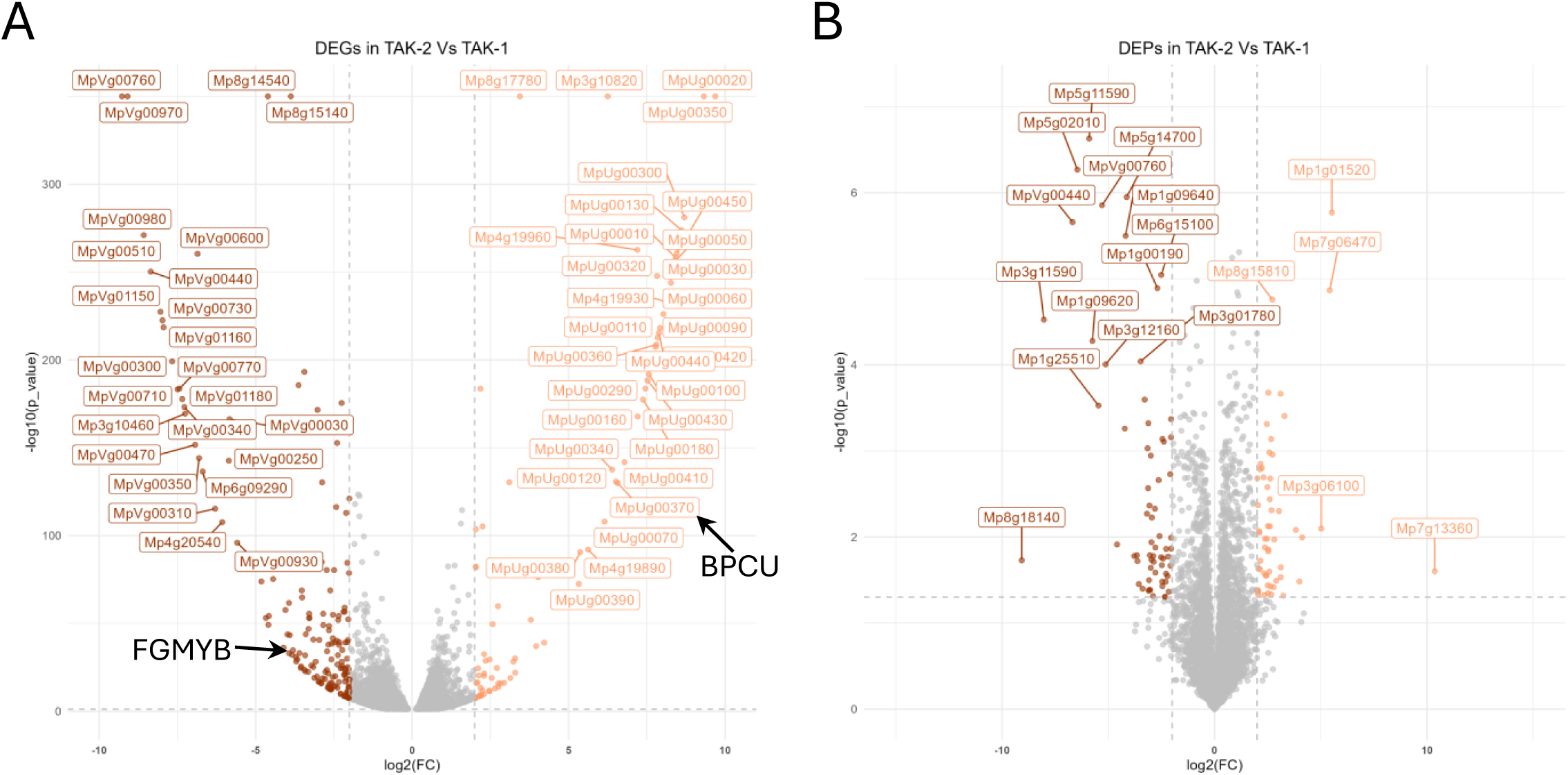
Volcano plots showing differential abundance of transcripts and proteins in the *Marchantia* genotypes Tak-1 (male) and Tak-2 (female). A) Transcriptional analysis showing that the majority of differentially expressed genes (DEGs) come from the female U chromosome and the male V chromosome. *BASIC PENTACYSTEINE ON THE U* (*BPCU*, MpUg00370) is the 24th most up-regulated gene in Tak-2 vs Tak-1, it suppresses expression of the autosomal transcription factor *FEMALE GAMETOPHYTE MYB* (*FGMYB*, Mp1g17210), the 40th most down-regulated gene of Tak-2 vs Tak-1. This is known to promote female sexual differentiation. B) Fewer differentially expressed proteins (DEPs) were identified by proteomics.

**Figure S2:**
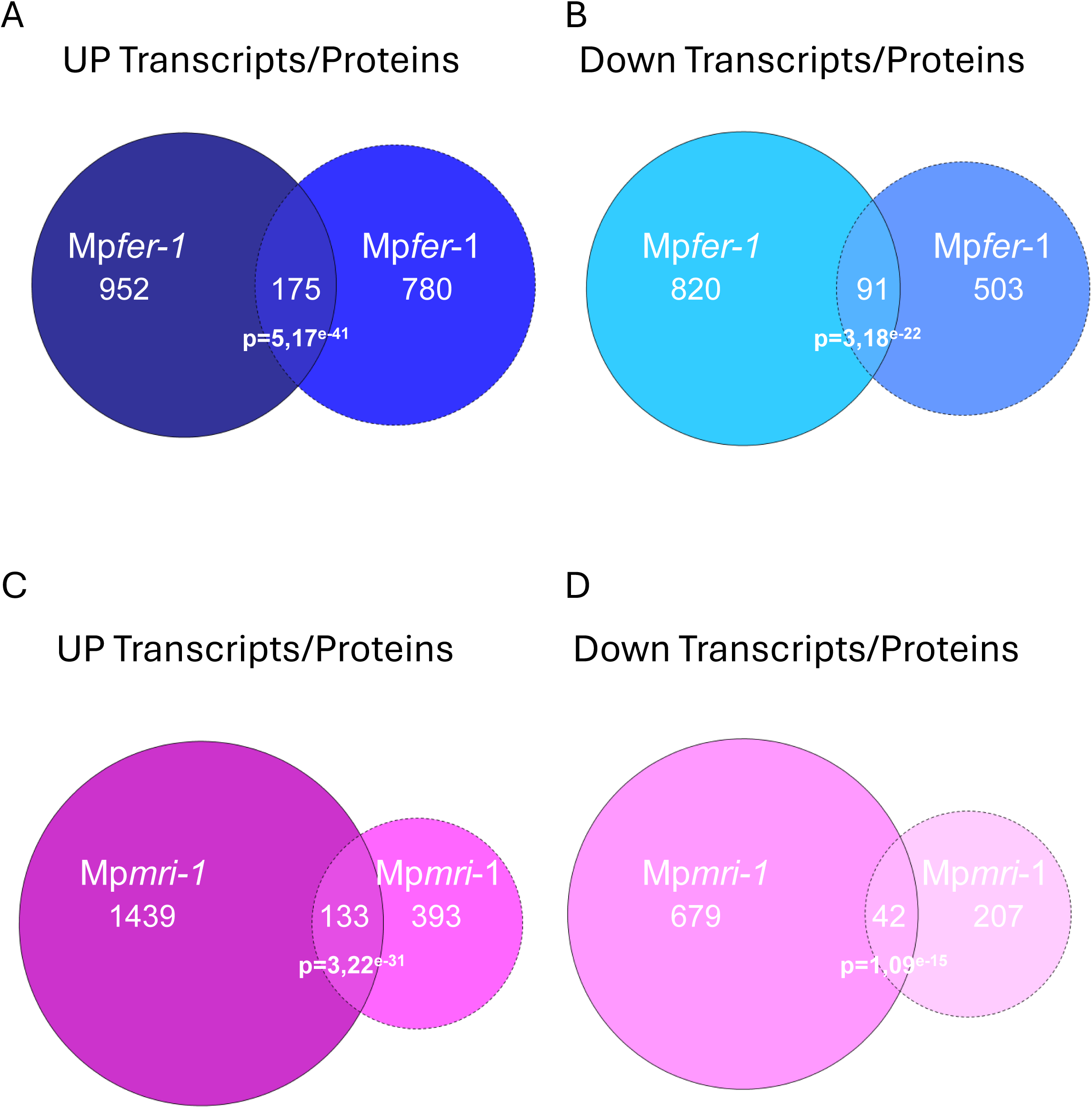
Venn diagrams showing the overlaps between differentially expressed transcripts and proteins in the *Marchantia* CWI mutants. A) Comparison of up-regulated DEGs and up regulated DEPs as well as B) down-regulated DEGs and DEPs for Mp*fer-1* vs Tak-1. C) Comparison of up-regulated DEGs and DEPs as well as (D) down-regulated DEGs and DEPs for Mp*mri-1* vs Tak-2. Statistical significance (p-values) of overlapping datasets was determined using a hypergeometric test.

**Figure S3:**
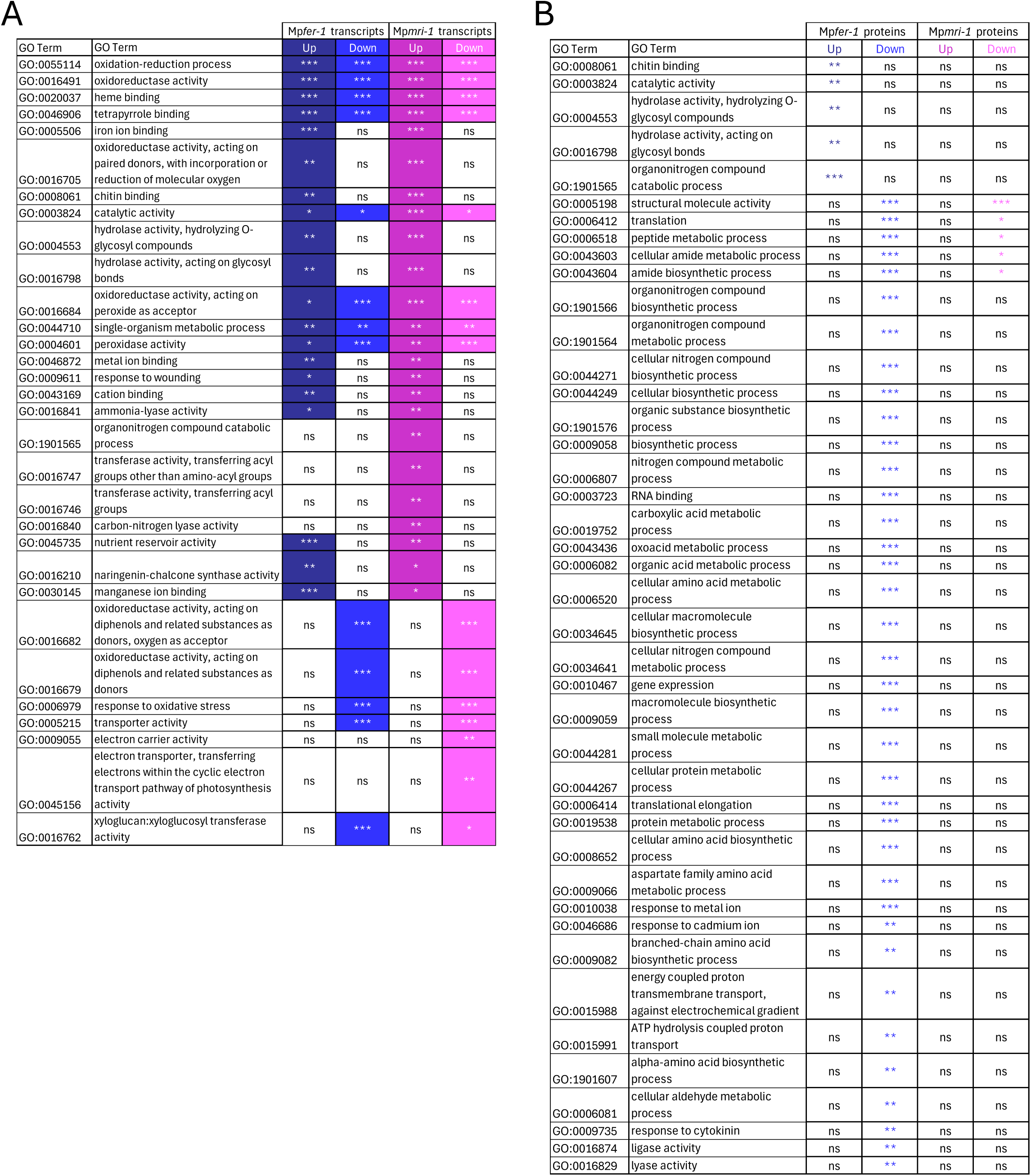
Gene ontology (GO) enrichment analysis in the *Marchantia* CWI mutants. A) Table showing GO categories significantly enriched in the transcriptomic data sets. B) Table showing GO categories significantly enriched in the proteomic data sets. Three stars (***) indicates statistical significance p≤0.001, two stars (**) indicates p≤0.01, one star (*) indicates p≤0.05, and ‘ns’ indicates not significant.

**Figure S4:**
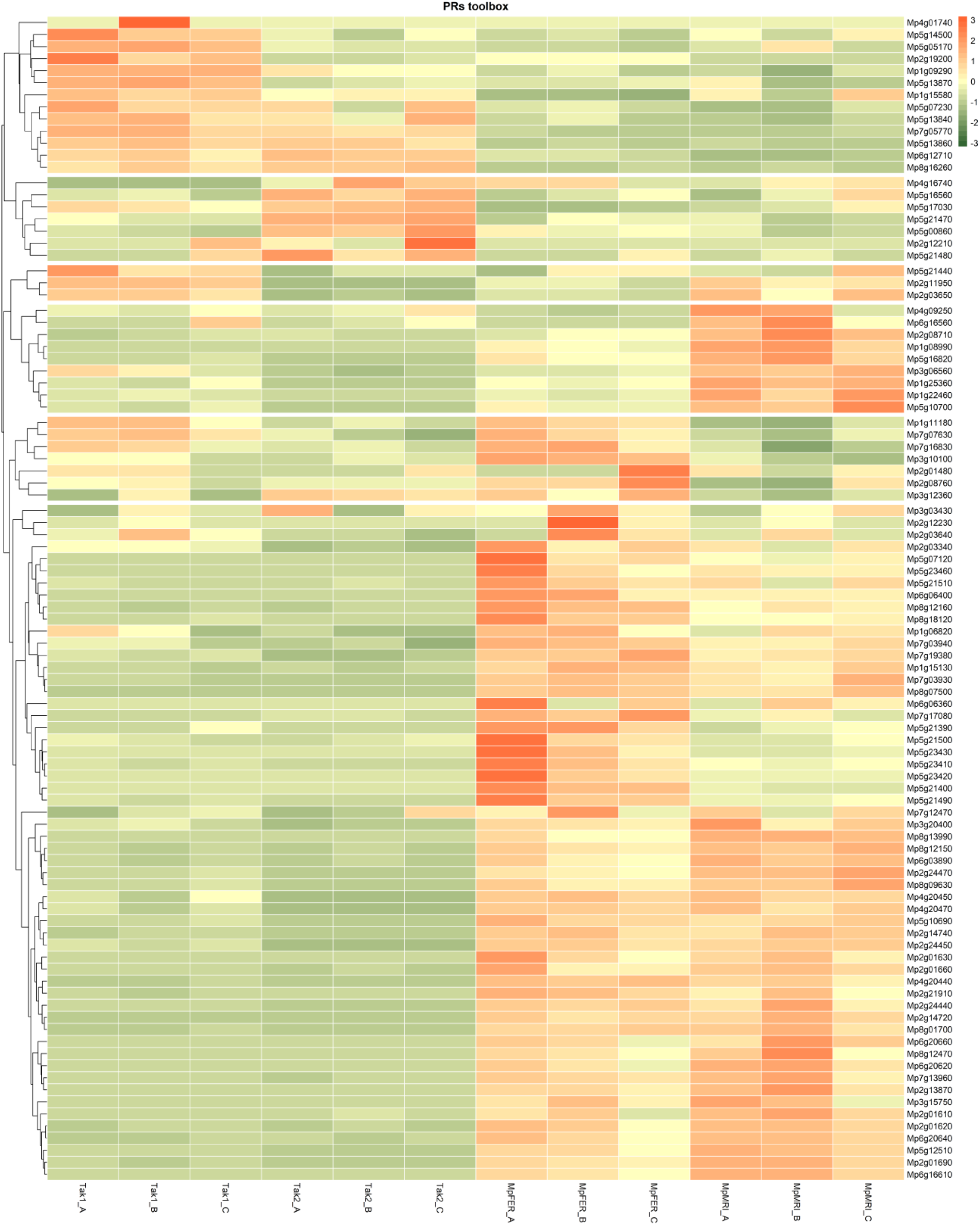
Heatmap and hierarchical clustering of *pathogenesis-related* (*PR*) gene expression across all biological replicates for Tak-1, Tak-2, Mp*fer-1*, and Mp*mri-1*. Each column represents a single biological replicate. Expression values were normalized by Z-score transformation.

**Figure S5:**
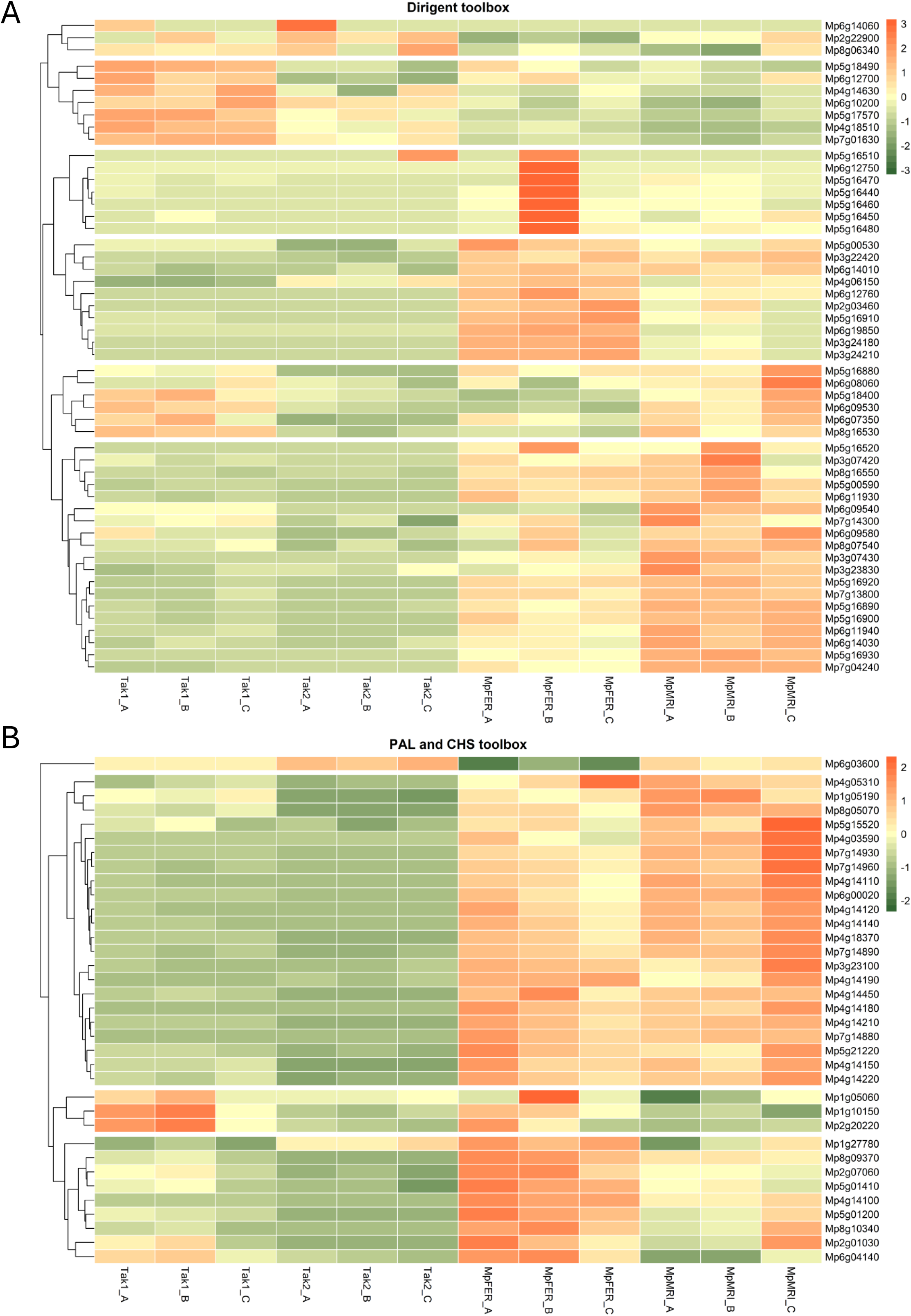
Heatmaps and hierarchical clusterings of *DIR* and *PAL/CHS* gene expression across all biological replicates for Tak-1, Tak-2, Mp*fer-1*, and Mp*mri-1*. A) *DIR* genes and B) *PAL/CHS* genes. Each column represents a single biological replicate. Expression values were normalized by Z-score transformation.

**Figure S6:**
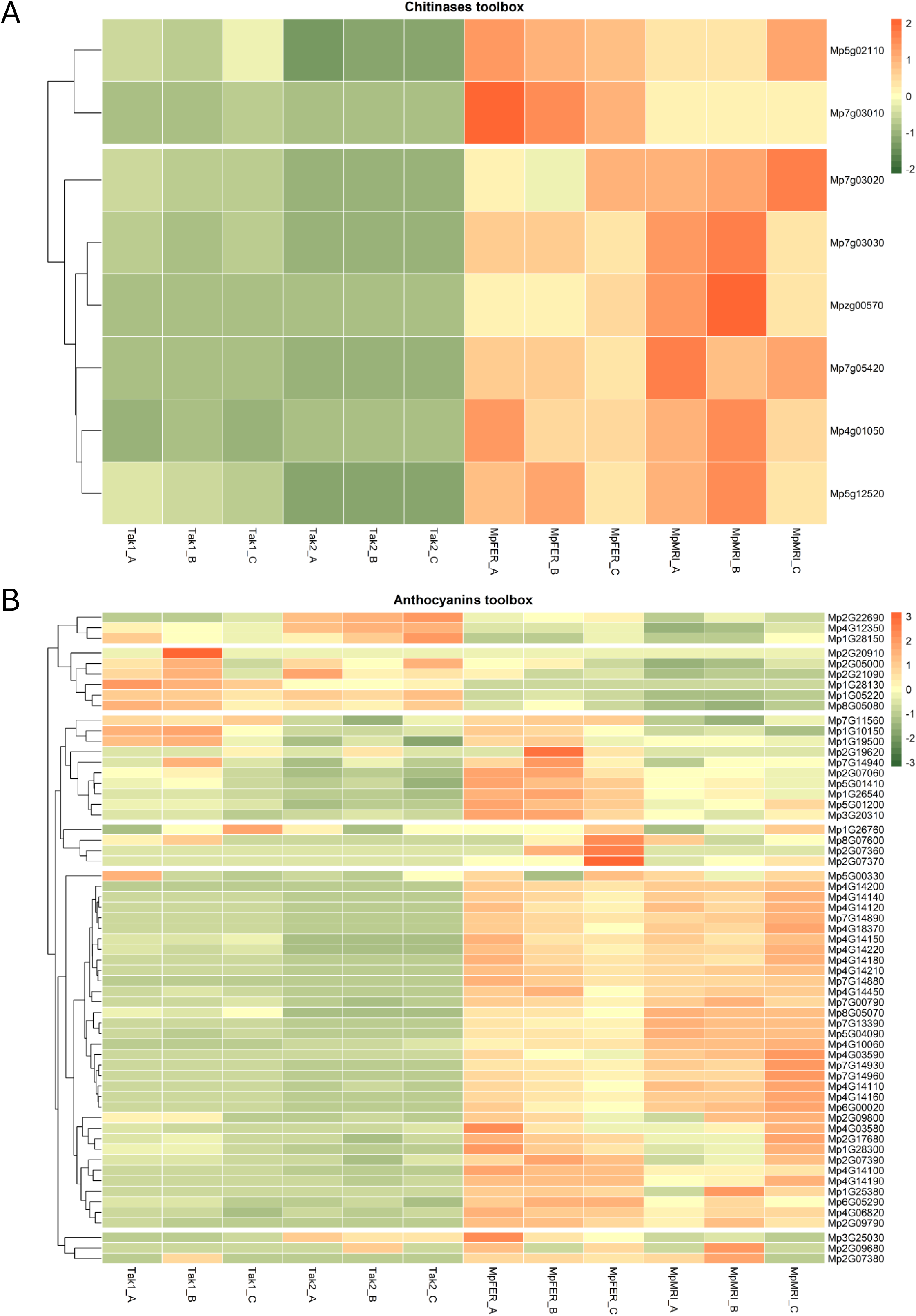
Heatmaps and hierarchical clusterings of *Chitinases* (A) and Anthocyanins-related (B) gene expression across all biological replicates for Tak-1, Tak-2, Mp*fer-1*, and Mp*mri-1*. Each column represents a single biological replicate. Expression values were normalized by Z-score transformation.

**Figure S7:**
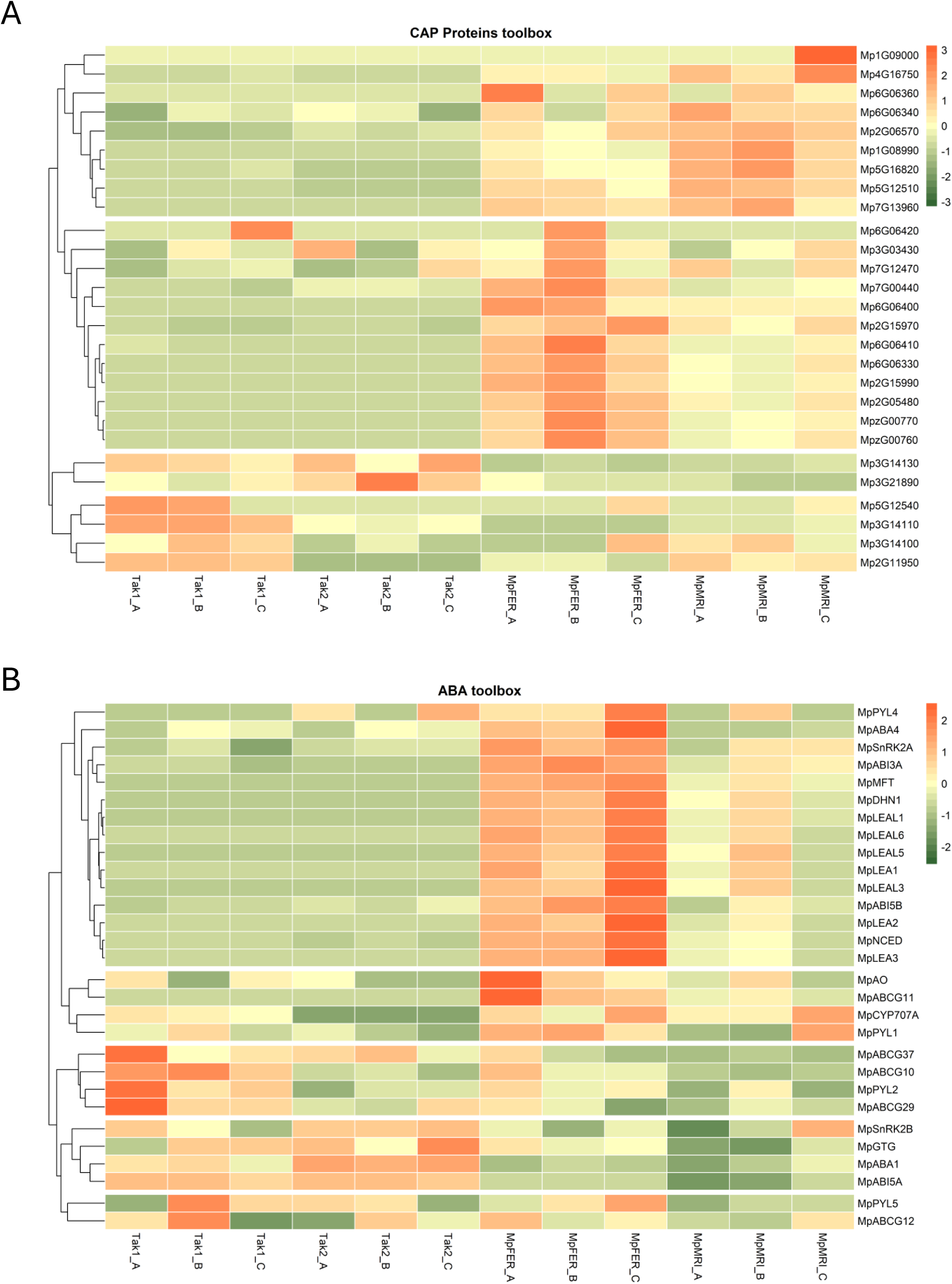
Heatmaps and hierarchical clusterings of *CAP* genes (A) and genes related to ABA (B) across all biological replicates for Tak-1, Tak-2, Mp*fer-1*, and Mp*mri-1.* Each column represents a single biological replicate. Expression values were normalized by Z-score transformation.

**Figure S8:**
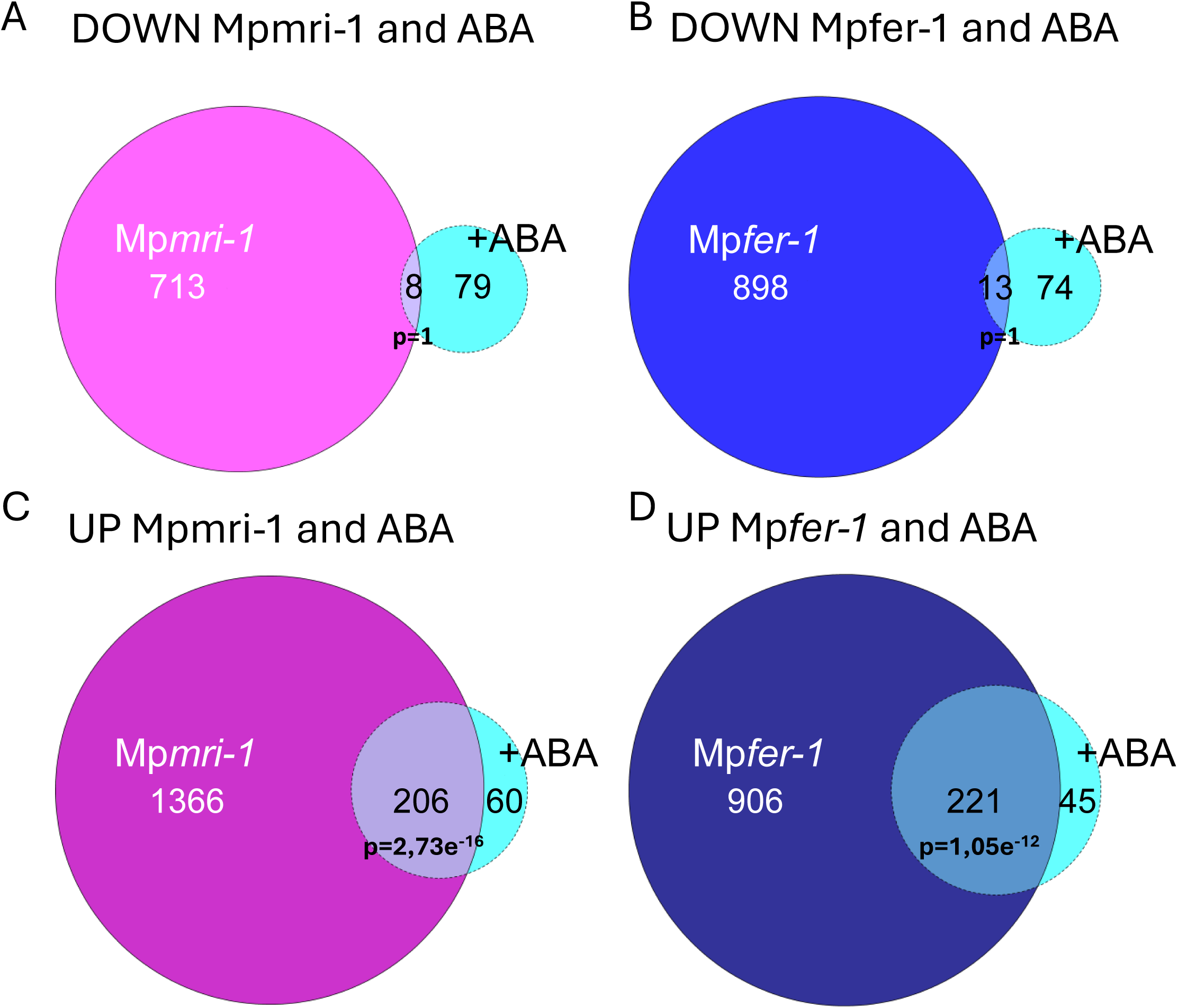
Venn diagrams showing differential abundance of transcripts in the untreated *Marchantia* CWI mutants compared to wild-type Tak-1 treated with 1 μM ABA. A) Transcriptional analysis showing DEGs that are down regulated in Mp*mri-1* compared with DEGs that are down regulated in WT under ABA treatment. B) Transcriptional analysis showing DEGs that are down in Mp*fer-1* compared with DEGs that are down regulated in WT under ABA treatment. C) Transcriptional analysis showing DEGs that are up regulated in Mp*mri-1* compared with DEGs that are up regulated in WT under ABA treatment. D) Transcriptional analysis showing DEGs that are up in Mp*fer-1* compared with DEGs that are up regulated in WT under ABA treatment. Differential expression was determined based on log_2_ (FC) ≥ 1.5 and p-value ≤ 0.05. The statistical significance of overlapping datasets was determined using a hypergeometric test.

**Figure S9:**
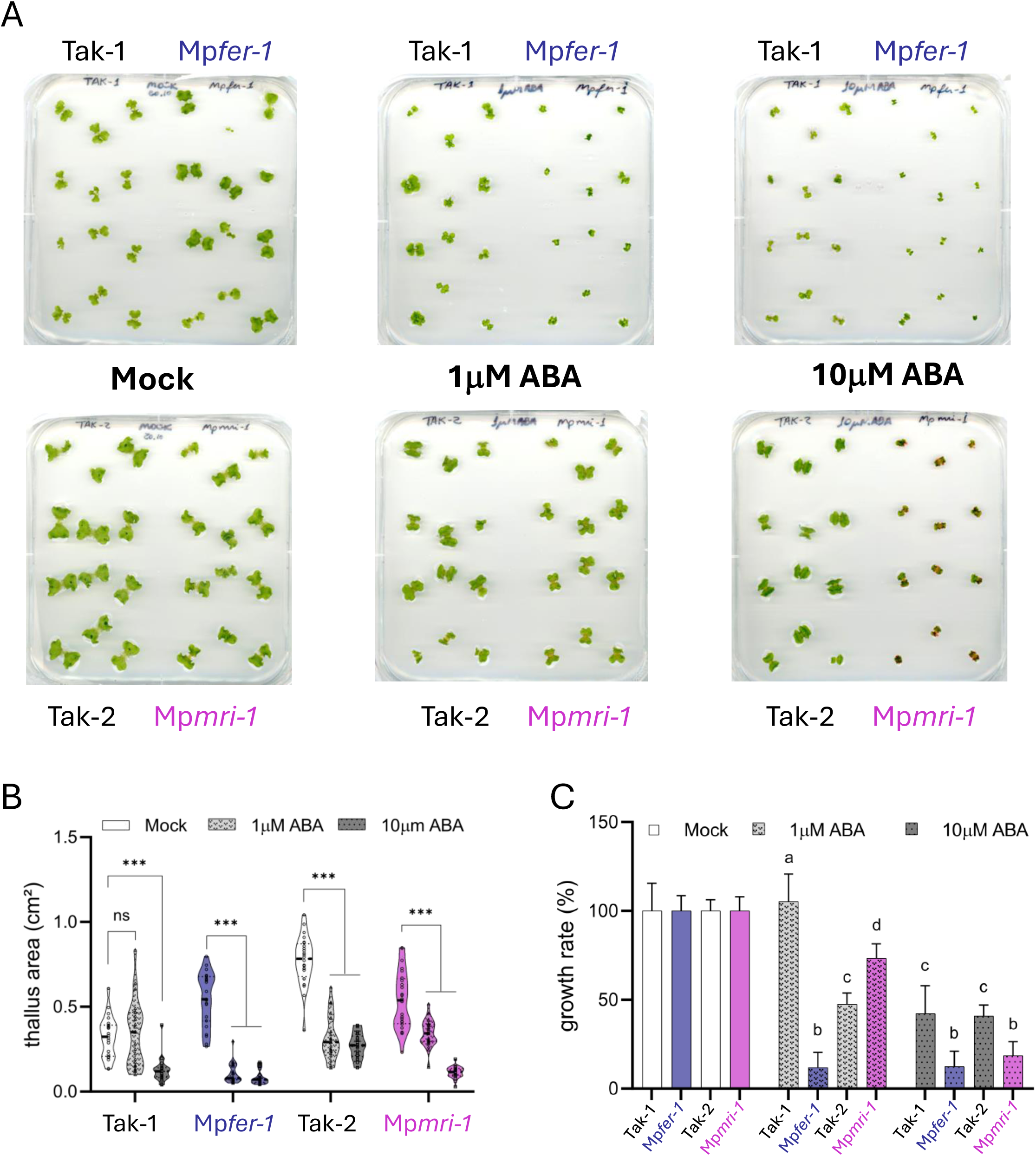
Independent assay of ABA treated Marchantia CWI mutants and quantification of thallus area and growth rate under ABA treatment. A) Representative images of *Marchantia* Tak-1, Tak-2, Mp*fer-1*, and Mp*mri-1* genotypes grown on agar plates supplemented with 0 μM ABA (mock), 1 μM ABA, or 10 μM ABA. B) Violin plot showing the thallus area of the four genotypes grown with 0 μM ABA (mock), 1 μM ABA, or 10 μM ABA. C) Bar chart showing the calculated growth rate as a fraction of the average of ABA-treated plant surface area to the average of Mock-treated surface area. Statistically significant differences were determined by two-way analysis of variance (ANOVA), followed by Tukey’s method with a significant level of P ≤0.05.

**Figure S10:**
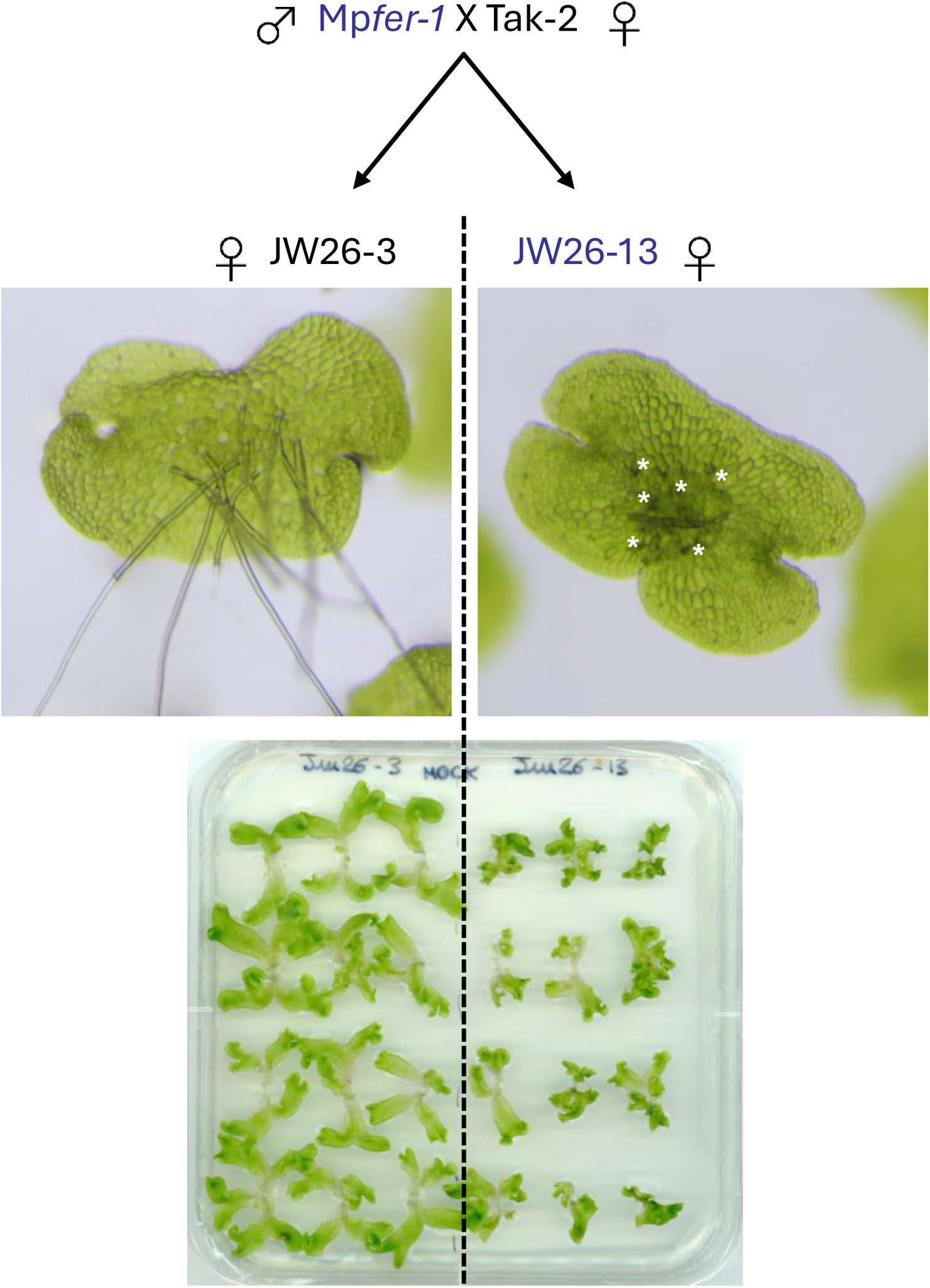
Mp*fer-1* (male) was crossed with Tak-2 (female). Among the progeny, 2 female individuals were recovered that segregated for the T-DNA at the Mp*FER* locus. JW26-3 did not harbor the T-DNA and displayed long intact rhizoids with normal thalli growth. In contrast, JW26-13 plants carried the T-DNA in Mp*FER*, displayed bursting short rhizoids (asterisks) and grew much smaller than JW26-3.

**Figure S11:**
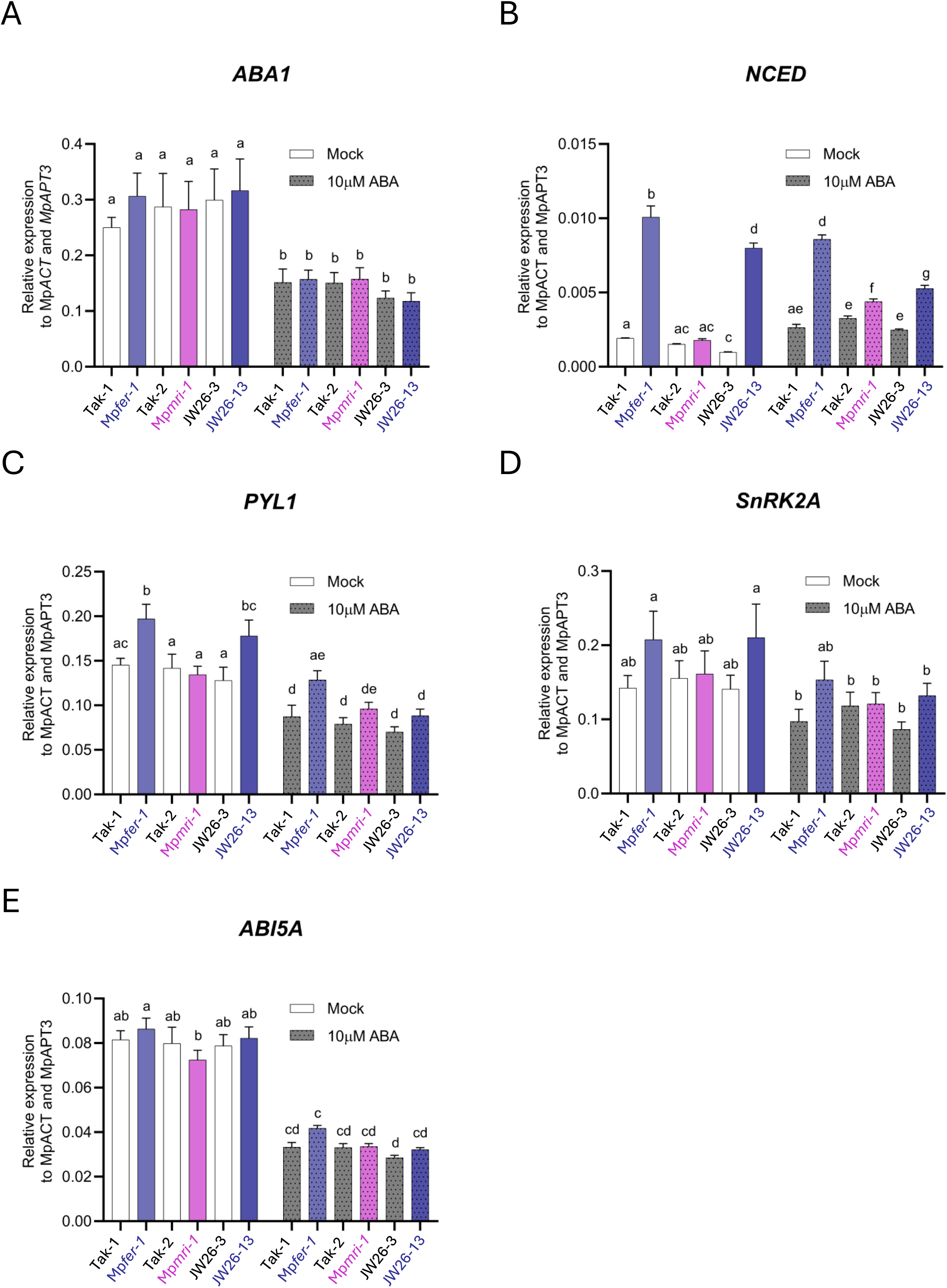
Gene expression analysis in Tak-1, Tak-2, Mp*fer-1,* Mp*mri-1,* JW26-3, and JW26-13 genotypes grown for 14 days on 10 μM ABA using quantitative real-time PCR (qRT-PCR). Transcripts levels for ABA-related genes were quantified, including the ABA biosynthesis genes A) Mp*ABA1* and B) Mp*NCED*, C) the ABA receptor Mp*PYL1*, the early ABA signaling gene D) Mp*SnRK2A*, and E) the ABA transcription factor Mp*ABI5A*. Normalization was performed against Mp*APT3* and Mp*ACT7*. Statistically significant differences were determined by two-way analysis of variance (ANOVA), followed by Tukey’s method with a significant level of P ≤0.05.

